# Massively parallel characterization of psychiatric disorder-associated and cell-type-specific regulatory elements in the developing human cortex

**DOI:** 10.1101/2023.02.15.528663

**Authors:** Chengyu Deng, Sean Whalen, Marilyn Steyert, Ryan Ziffra, Pawel F. Przytycki, Fumitaka Inoue, Daniela A. Pereira, Davide Capauto, Scott Norton, Flora M. Vaccarino, Alex Pollen, Tomasz J. Nowakowski, Nadav Ahituv, Katherine S. Pollard

## Abstract

Nucleotide changes in gene regulatory elements are important determinants of neuronal development and disease. Using massively parallel reporter assays in primary human cells from mid-gestation cortex and cerebral organoids, we interrogated the *cis*-regulatory activity of 102,767 sequences, including differentially accessible cell-type specific regions in the developing cortex and single-nucleotide variants associated with psychiatric disorders. In primary cells, we identified 46,802 active enhancer sequences and 164 disorder-associated variants that significantly alter enhancer activity. Activity was comparable in organoids and primary cells, suggesting that organoids provide an adequate model for the developing cortex. Using deep learning, we decoded the sequence basis and upstream regulators of enhancer activity. This work establishes a comprehensive catalog of functional gene regulatory elements and variants in human neuronal development.

**One Sentence Summary:** We identify 46,802 enhancers and 164 psychiatric disorder variants with regulatory effects in the developing cortex and organoids.

## Introduction

Psychiatric disorders affect nearly one in five adolescents in the United States (*1*) and have a strong genetic etiology (*2*). Studies profiling gene expression across distinct anatomical regions have found an enrichment for psychiatric disorder-associated genes in stages of developmental neurogenesis in the marginal zone and deep cortical layer neurons (*2*–*4*). For example, most autism spectrum disorder (ASD) risk genes were found to regulate gene expression or be involved in neuronal communication during early brain development (*5*). Analysis of genes associated with schizophrenia via genome-wide association studies (GWAS) found several to be expressed in the prefrontal cortex at early developmental stages (*6*). Thus, decoding the genetic causes of psychiatric disorders requires deep knowledge of gene regulatory mechanisms in the developing brain.

In the past decade, hundreds of psychiatric disorder-associated genetic risk loci have been identified, both by work of individual labs and large consortia, including the Psychiatric Genomics Consortium (*7*, *8*) and psychENCODE (*9*). A major portion of these loci reside in noncoding regions of the genome, likely within gene regulatory elements, and contain highly correlated variants due to linkage disequilibrium (LD), making them challenging to interpret and functionally characterize. Gene regulatory elements, such as enhancers and promoters, are known to regulate lineage- and region-specific transcription in the developing human cortex (*10*). While promoters are located adjacent to their target genes, enhancers can be located at distal locations from the gene(s) that they regulate. In addition, due to their cell-type specificity and spatiotemporal dynamic activity, enhancers are difficult to identify. Single-cell ATAC-seq (scATAC-seq) at different developmental stages of the human cortex enabled the identification of different cell populations and their candidate regulatory elements in the developing cortex (*11*, *12*). This work also showed significant correlations between these cell types and cerebral organoids. However, these studies are descriptive and do not provide a functional readout that can test the enhancer activity of these sequences and the effects of psychiatric disorder-associated variants on their function.

Massively parallel reporter assays (MPRAs) enable the simultaneous testing of thousands of sequences for their regulatory activity (*13*). The quantitative readout of an MPRA makes it possible to test different alleles of the same locus side-by-side in one experiment, enabling the detection of sequence variants that alter enhancer function (*14*–*16*). Because they generate data for large numbers of sequences, MPRAs have enabled the development of deep learning models that predict enhancers and their quantitative activity from sequence alone (*17*, *18*). Machine learning has been used to predict: 1) enhancers from epigenetic marks, sequence motifs, and evolutionary conservation (reviewed in (*19*)); 2) MPRA activity from sequences and epigenetic features (*20*); 3) gene expression from promoter-proximal sequences (*21*–*23*); and 4) epigenetic features from sequences (*24*–*30*). In addition, DNA sequence-based models (*17*, *18*) have the potential to be deployed at scale to screen variants without MPRA data and to design enhancers with desired properties, such as cell type specificity. These strategies have shed light on the sequence motifs and upstream regulators that are most important for regulating gene expression across different cell types and species.

Here, we used deep learning and a lentivirus-based MPRA (lentiMPRA) to test 102,767 sequences for their enhancer activity in primary human mid-gestation cortical cells and 10-week cerebral organoids. These sequences included cell-type-specific open chromatin regions and psychiatric disorder-associated quantitative trait loci (QTLs) predicted to be functional enhancers based on their epigenetic profiles. Combined, we discovered 46,802 sequences to be functional enhancers and 164 variants with significant allelic differences in enhancer activity regulating known disorder-associated genes such as *TBR1*, *MARK2* (autism spectrum disorder), and *NFKB2* (schizophrenia). We observed comparable activity levels between organoids and primary cortical cells, suggesting that organoids provide an adequate *in vitro* model to study the developing cortical regulatory landscape. Finally, we used our lentiMPRA data to train a deep learning model that predicts enhancer activity from sequence with state-of-the-art accuracy, enabling us to learn sequence determinants and upstream regulators of the human mid-gestation enhancer code. These findings provide a comprehensive catalog of functional cortical-cell-specific enhancers and psychiatric disorder-associated variants that alter their activity, improving our understanding of the molecular basis of neurodevelopment and the etiology of psychiatric disorders.

## Results

### LentiMPRA library generation and analysis

To comprehensively characterize human neurodevelopmental enhancers and their sequence variants across the mid-gestation cortex, we designed two lentiMPRA libraries (Methods) and tested them in primary human cortical cells (**Fig. 1A**). Due to the limited number of obtainable human primary cells and lentivirus integrations into these cells, each library was assayed independently.

**Fig. 1.**
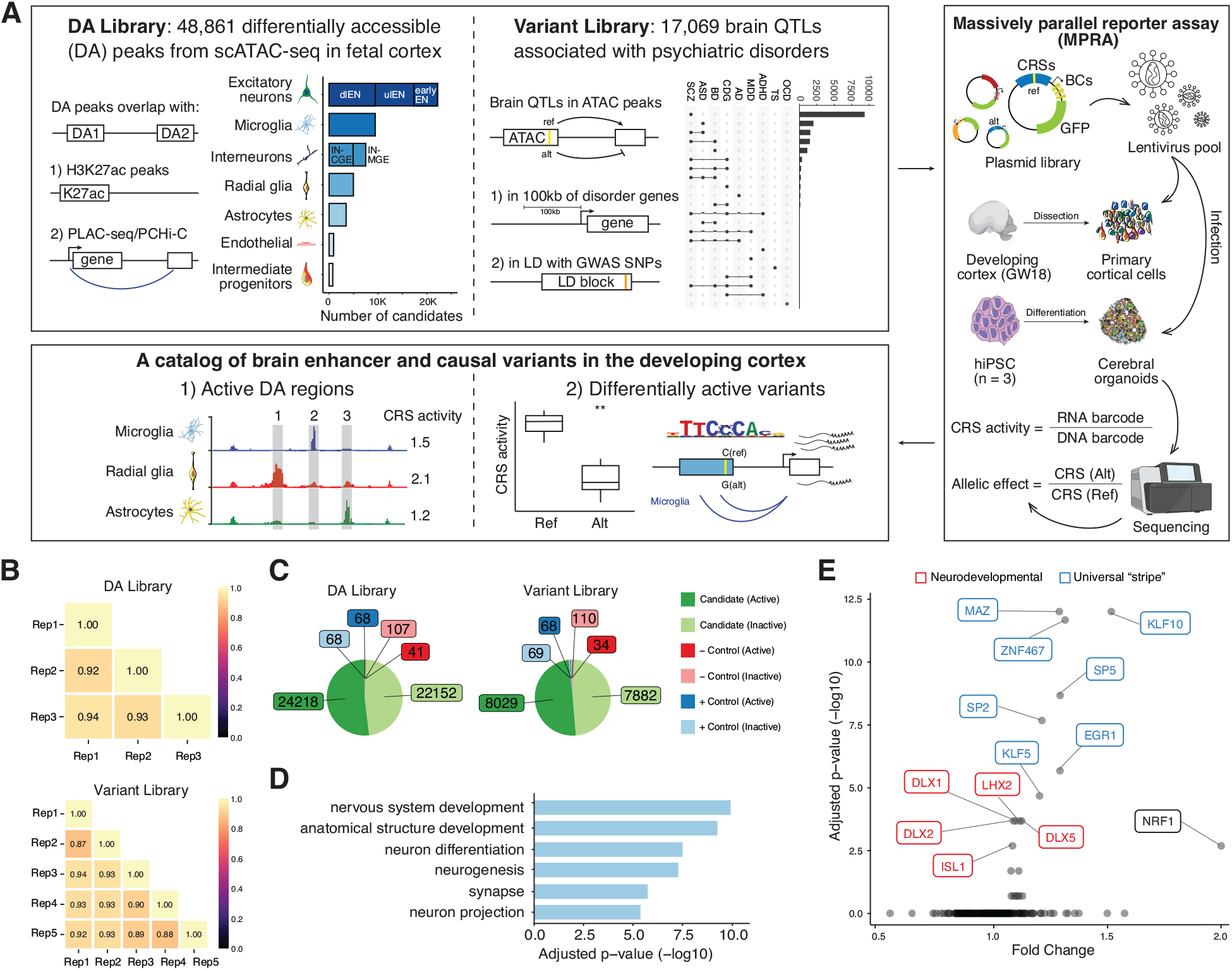
Design and overall lentiMPRA results. (**A**) Experimental overview of the two lentiMPRA libraries. Library 1 contains 48,861 differentially accessible (DA) regions from scATAC-seq in the developing human cortex that either overlap H3K27ac peaks or PLAC-seq/PCHiC loop. The number of DA candidates for each cell type was portrayed in the bar plot. dlEN, deep layer excitatory neuron; ulEN, upper layer excitatory neuron; IN-CGE, CGE derived interneuron; IN-MGE, MGE derived interneuron. Library 2 includes 17,069 brain QTLs that are 100kb from differentially expressed cross-disorder neurodevelopmental genes or in linkage disequilibrium (LD) with psychiatric disorder GWAS SNPs. The number of variants associated with each disorder is shown in the upset plot (top 22 intersects shown). SCZ, schizophrenia; ASD, autism spectrum disorder; BD, bipolar disorder; CDG, congenital disorders of glycosylation; AD, Alzheimer’s disease; MDD, major depressive disorder; ADHA, attention-deficit/hyperactivity disorder; TS, Tourette syndrome; OCD, obsessive-compulsive disorder. Both libraries were cloned into a lentiMPRA vector and packaged into lentivirus and used to infect primary cortical cells dissociated from GW18 tissues and human induced pluripotent cell (hiPSC)-derived cerebral organoids. Following infection, DNA and RNA were extracted and sequenced and an RNA/DNA barcode count ratio was calculated for each candidate regulatory sequence (CRS) allowing the identification of active DA regions and differentially active variants. (**B**) Correlation of log2(RNA/DNA) between technical replicates in primary cortical cells for library 1 and 2, respectively. (**C**) Pie charts showing the number of active and inactive sequences for candidates, positive (+) and negative (−) controls in both libraries. (**D**) Top enriched GO terms from the ‘Biological Process’, ‘Cellular Component’ and ‘Molecular Function’ ontologies for nearest genes of the highest activity sequences (both libraries combined). Closest genes of the lowest activity sequences were used as the background set. The complete list of GO terms is available in fig. S2B. (**E**) TF motif enrichment analysis for highest activity sequences (both libraries). Red: neurodevelopmental TFs, Blue: USFs.

The first library was designed to characterize cell-type specific enhancers. It consisted of 51,495 sequences obtained primarily from differentially accessible (DA) scATAC-seq peaks in the developing human brain (*11*). These DA peaks were further selected based on their: 1) overlap with histone H3 acetylation of lysine 27 (H3K27ac) peaks from bulk prefrontal cortex (PFC) tissue (*31*), microglia or non-microglia cells (*11*) (n = 24,611, 53%); or 2) overlap with H3K4me3 proximity ligation-assisted chromatin immunoprecipitation sequencing (PLAC-seq) peaks from intermediate progenitor cells (IPC), radial glia (RG), excitatory neurons (EN), or interneurons (IN) (*32*) (n = 12,412, 26.8%); or 3) overlap with promoter capture Hi-C (PCHi-C) from EN, hippocampal dentate gyrus (GE)-like neurons, lower motor neurons and astrocytes (*33*) (n = 13,712, 29.5%).

The second library compared the reference and the alternative alleles for 17,069 variants. These were designed from brain QTLs (*34*–*36*) overlapping pseudo-bulked ATAC-seq peaks (*11*) that: 1) are within 100 kilobases (kb) of differentially expressed genes in schizophrenia, autism or bipolar disorder (expression QTL (eQTL) n = 14,021; chromatin QTL (caQTL) n = 149) (*9*); or 2)share a linkage disequilibrium (LD) block with GWAS SNPs for various psychiatric disorders (*8*, *37*–*44*) (eQTL n = 2,882, cQTL n = 17). Assaying both QTL and GWAS SNPs allowed us to overcome the systematic bias toward different types of variants in each type of study (*45*), thus enabling a comprehensive screening for functional regulatory variants. This library also included ~15,000 non-QTL sequences with a range of expected activity levels predicted from their epigenetic profiles.

To prioritize distal enhancers, promoter-overlapping peaks were excluded from both libraries. Each library contained 143 positive control sequences nominated from ATAC-seq and ChIP-seq data in brain organoid models (*46*) and used to define active sequences, plus ~2,000 additional controls used for quality control. We synthesized 270 base pair (bp) oligos, each centered on either the DA peak summit (library 1) or variant (library 2) followed by 15bp primers on either side to amplify the library. A 31bp minimal promoter (minP) and 15bp random barcode were placed downstream of each synthesized oligo via PCR and cloned into a lentiMPRA vector (**Fig. 1A**).

Both libraries were packaged into lentivirus and used to infect human primary cortical cells at mid-gestational week (GW) 18. Primary cortical cells were cultured for two days before infection, exhibiting complex morphology and smooth neurites. The presence of major cortical cell types was confirmed by immunocytochemistry before and after infection, including deep-layer excitatory neurons (dlENs), upper-layer excitatory neurons (ulENs), newborn excitatory neurons (earlyENs), RG, IPC, astrocyte/oligodendrocyte precursors (Astro/oligo), endothelial and mural cells (EndoMural), medial ganglionic eminence derived interneurons (IN-MGE), caudal ganglionic eminence derived interneurons (IN-CGE), and microglia (MG) (**fig. S1A**). We performed three replicates for the DA library and five replicates for the variant library. Three days post-infection, when the majority of the non-integrated virus was gone, DNA and RNA were harvested and prepared for sequencing. DNA sequencing revealed that both libraries contained more than 96% of the designed oligos (DA library: 50,394 oligos; variant library: 51,319 oligos), and each oligo had on average over 50 unique barcode associations (median DA: 56, variant: 64). Overall, 97,762 sequences (95%) passed stringent quality control (Methods).

To measure enhancer activity, we quantified depth-normalized barcode abundance in DNA and RNA for each oligo and then calculated its batch-corrected RNA/DNA ratio. A high correlation of RNA/DNA ratios between replicates was observed (average Pearson correlation, DA: 0.93, variant: 0.91; **Fig. 1B**), confirming sufficient reproducibility. We next compared the activity distributions of positive and negative controls (**fig. S2A**). As expected, positive controls had significantly higher ratios than negative controls in both libraries (DA: p = 1e-3, variant: p = 8e-5, Wilcoxon test). Moreover, the distribution of ratios for randomly scrambled negative controls was highly comparable between libraries (median DA: 0.997, median variant: 0.994), indicating that activity was mostly driven by the minimal promoter. Altogether, these quality assessments suggest that our lentiMPRA robustly distinguishes between sequences with high, medium, and low regulatory activity.

To identify sequences capable of driving gene expression, we defined active sequences as those with RNA/DNA ratios higher than the median of positive controls in their respective libraries (DA: 1.047, variant:1.068), conservatively treating the remaining sequences as inactive. We further defined the highest activity sequences as those above the 75th percentile of the positive controls and the lowest activity sequences as those below the 25th percentile. Combining both libraries, we identified a total of 46,802 active sequences (48% of 97,762) and 25,557 with the highest activity. We next evaluated the various properties of active lentiMPRA sequences. Compared to inactive sequences, active sequences are significantly more conserved (p=5.8e-28, Wilcoxon test), and their target genes are expressed at higher levels during mid-gestation (p=6.4e-6, Wilcoxon test; 23% of all sequences mapped to target genes using PLAC-seq data (*32*)). Comparing the highest versus lowest activity sequences, we found gene ontology (GO) enrichment for neurodevelopmental terms, such as ‘nervous system development’ and ‘neuron differentiation’ (**Fig. 1D**). Next, we analyzed transcription factor binding sites (TFBSs) and observed enrichment for known neurodevelopmental TFBSs including the *DLX*, *LHX* and *SOX* gene families. We also found enrichment for universal stripe factors (USFs) including *EGR1*, *MAZ* and members of the *KLF/SP* family (**Fig. 1E**). USFs colocalize at most promoters and enhancers, increasing chromatin accessibility and residence time for cofactors (*47*), and our finding suggests that they play a similar role with lentiMPRA reporter constructs integrated into the genome. Together, these results indicate that our active sequences have biological functions in brain development.

### Identification of thousands of active cortical-cell-type specific enhancers

Of 46,370 DA sequences passing quality control, 24,218 (52%) were active enhancers in primary cortical cells (**Data S1**). Among these active DA, 4,656 (19%) were transiently accessible regions across pseudotime of excitatory neurogenesis in a previous scATAC-seq study (*11*). We then separated DA sequences based on their cell-type specificity and found that for each cell type 43-62% of the sequences were active (**Fig. 2A**). We observed that abundant cell types, including neurons and radial glia, had a slightly higher percentage of active sequences compared to less abundant cell types, such as microglia and endothelial/mural cells. Compared to the lowest activity DA sequences, the highest activity DA sequences are enriched for TFBS of neurodevelopmental transcription factors, and many of these show positional enrichment within the ATAC-seq peak (**Fig. 2B**). For example, active sequences tend to have motifs for ATOH1, NEUROD2, and TCF4 upstream of the peak summit, whereas ASCL1 and SPI1 motifs are enriched downstream of the summit.

**Fig. 2.**
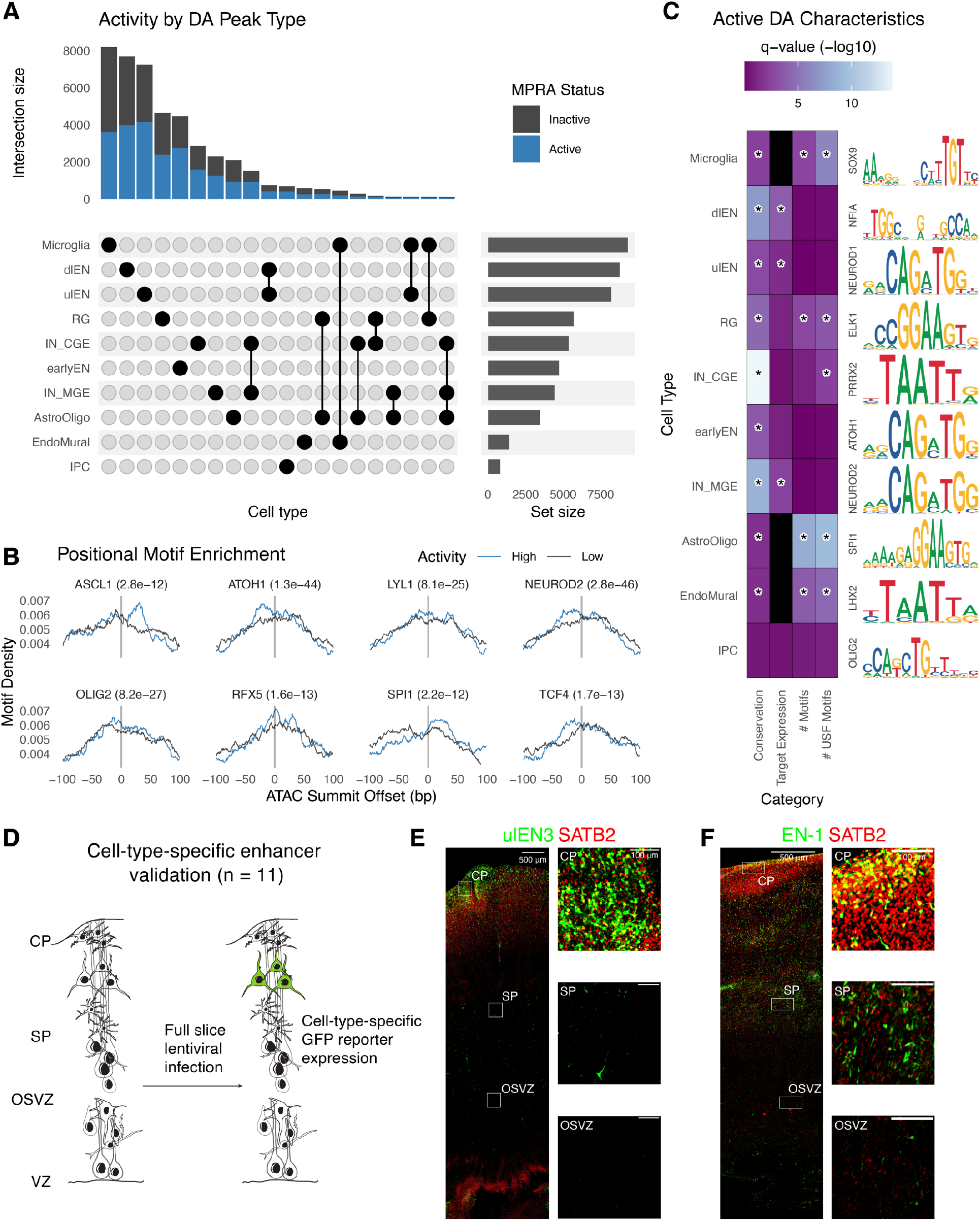
Identification and validation of functional differentially accessible regions in the developing cortex. (**A**) Upset plot showing the number of DA peaks (active: blue; inactive: gray) for each cell type or combination of cell types. (**B**) The highest activity DA sequences have positional motif enrichment for neurodevelopmental TFs compared to the lowest activity sequences, exhibiting significantly more motif matches slightly up- or downstream of the ATAC-seq peak summit. (**C**) Active DA sequences have significantly higher means across several attributes compared to inactive DA sequences (color scale: Wilcoxon test q-values, black = no data) including evolutionary conservation (phyloP), expression of PLAC-seq linked target genes in matched cell types, total number of strong motif matches (q-value < 0.01), and total number of strong USF motif matches. A representative motif enriched in active DA peaks for each cell type is shown on the right. Statistically significant comparisons (q-value < 0.05) are indicated by a star. (**D**) Experimental strategy for validating cell type specificity of active DA sequences. (**E-F**) Mid-gestation human cortex slice cultures transduced with a GFP lentivirus reporter driven by a ulEN-specific enhancer (chr5:89274678-89274948, hg38) (**E**) and a pan-excitatory neuron specific enhancer (chr2:165141999-165142269, hg38) (**F**). Expression of GFP (green) and *SATB2* (red) was visualized via immunohistochemistry staining and insets show colocalization of GFP+ along with *SATB2*+ cells in different layers.

Next, we assessed the characteristics of active DA sequences across cell types (**Fig. 2C and table S1**). In almost all cell types, active DA sequences are more conserved than inactive sequences, consistent with prior knowledge that neurodevelopmental enhancers tend to exhibit strong conservation across vertebrate evolution (*48*). Inhibitory neurons derived from the ganglionic eminence exhibited the largest differences in conservation scores between active and inactive sequences, fitting with the general transitory role of the ganglionic eminence in guiding neuronal migration (*49*). Next, to test whether active DAs had regulatory activities endogenously, we predicted the target genes of each DA sequence using PLAC-seq data (*32*) and calculated cell-type-matched expression using scRNA-seq data in the developing human cortex (*11*). For many neuronal subtypes, genes interacting with active DAs showed higher expression compared to genes interacting with inactive DAs. For example, putative genes regulated by dIEN-specific active DAs were transcribed at a higher level in dIEN, compared to genes regulated by dlEN-specific inactive DAs (p = 0.00016, Wilcoxon test). We also found a significant higher number of strong TF motifs in active DAs specific to AstroOligo, EndoMural, Microglia and RG, while USF motifs were enriched in DAs specific to IN-CGE plus these same four glial and vascular cell types. These results indicate that the activity of DA sequences in lentiMPRA is associated with motif content and target gene expression in the matched cell type. Lastly, we observed unique sets of enriched motifs in active DAs of each cell type, and found many of them formed functional protein-protein interaction (PPI) networks (**fig. S4**).

To further verify the cell-type-specific activity of active DAs in our lentiMPRA, we selected eleven DA peaks with high MPRA activity from six different cell types (**table S2**) and tested them individually for their enhancer function in cortical cells followed by co-staining with antibodies for cell-specific markers (**Fig. 2D**). Each sequence was individually cloned into our lentiMPRA vector in front of a GFP reporter construct. As tissue samples are difficult to obtain at exactly GW18, we used cortical sections covering the mid-gestation period more generally (GW16-20) for infection. We first validated that the cortical sections can be uniformly infected by transducing them with a constitutively active enhancer vector, finding that our virus was able to infect with even spatial distribution across cell types (**fig. S3**). We next infected tissues with the individual enhancer viruses and found that all candidates showed GFP expression (**fig. S3**), in agreement with our lentiMPRA results. Cell-type specificity was inferred from GFP spatial location, counterstaining with cell markers, and morphology. While some candidates did not display strong cell-type-specific signals, we found three excitatory neuron-specific DA sequences (EN-1, ulEN-2, and dlEN-2) showing enhancer activity in the expected cell type. For example, a ulEN-specific DA region (chr5:89274678-89274948, hg38, **Fig. 2E**) drove GFP expression predominantly in the upper areas of the cortical plate and largely co-localized with *SATB2*, an upper layer excitatory neuron marker. Using PLAC-seq data (*32*), we found that this region has an EN-specific interaction with the promoter of *MEF2C* and *MEF2C-AS1*, known ASD and SCZ genes with EN-specific expression in the developing cortex. Another example is a pan-excitatory neuron specific accessible region (chr2:165141999-165142269, hg38, **Fig. 2F**) that showed notably higher GFP signal in the cortical plate (CP) and subplate (SP) compared to the ventricular zone (VZ) and outer subventricular zone (OSVZ), with most of the cells positive for GFP and *SATB2* located in the top layer of CP.

Not all sequences showed enhancer activity within or unique to their predicted cell type. Two regions (ulEN1 and dlEN1) showed GFP expression outside the regions where the expected cell type is enriched and did not overlap with cell marker staining. Candidate sequences specific to Astro/Oligo or RG showed GFP signal around the VZ but also near the CP. In these cases, GFP+ cells showed complex morphologies: some matched with the expected cell type(s) while others did not (**fig. S3**). To conclude, we independently validated the enhancer activities of eleven sequences with high lentiMPRA activity, finding all of them to drive GFP expression in cortical cells, with three exhibiting cell-type-specificity consistent with predicted activity from scATAC-seq.

### lentiMPRA identifies functional regulatory variants associated with psychiatric disorders

We next characterized the psychiatric disorder-associated variant library. Of the 15,335 variants with both alleles passing quality control, 8,029 (52%) showed enhancer activity from at least one allele. For these active variants, we estimated the allelic effect on enhancer activity by testing for differential activity across replicates. Most variants had modest effect sizes (median absolute log_2_ FC = 0.069) (**Fig. 3A**). This is in line with previous MPRAs studying the effect of regulatory variants (*50*, *51*), and consistent with eQTL analyses that find ~1% of single nucleotide changes are associated with significant changes in gene expression (*52*). At a 10% false discovery rate, 164 variants showed significant allelic effects with the number of down-regulating and up-regulating variants being similar (51% versus 49%, p = 0.81, Binomial test, **Fig. 3B**) (*53*). Our subsequent analyses focus on these 164 differentially active variants (DAVs) (**Fig. 3B**, **Data S2**). Among DAVs, 26 were in LD with GWAS SNPs and 138 were within 100kb of differentially expressed disease genes, which is similar to expectation given the library design (17% GWAS and 83% eQTL). Consistent with being QTLs, DAVs are not enriched for low-frequency variants (OR=0.8, p=0.34) nor do they have elevated conservation (OR=0.88, p=0.52). Separating DAVs based on cell type showed enrichment in Astro/Oligo (OR=2.39, p=0.14, **Fig. 3C**), which is particularly striking given the relatively low proportion of this cell type in our cultures and indicates the importance of astrocytes and oligodendrocytes in psychiatric disorders.

**Fig. 3.**
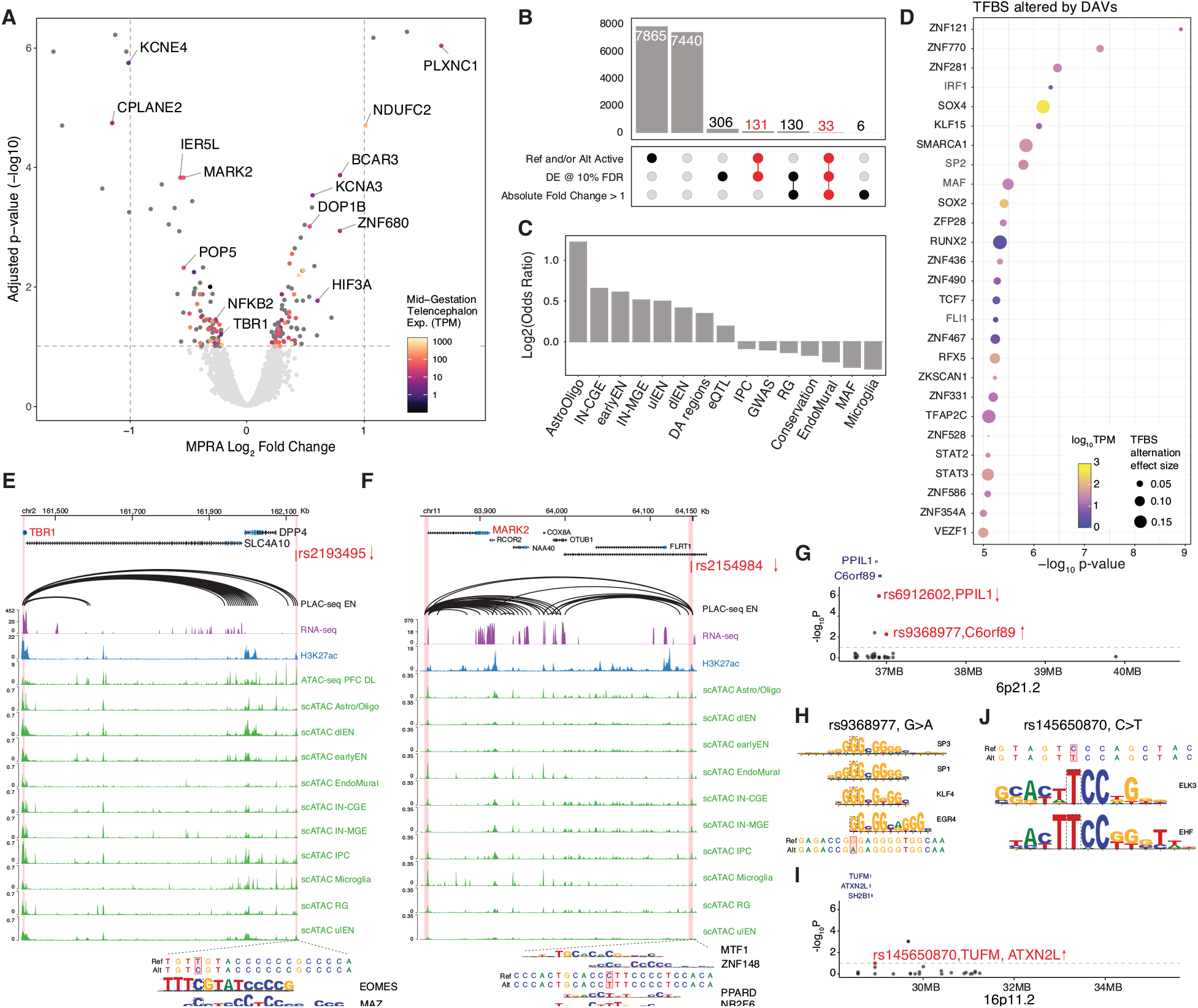
Identification of differential active variants associated with psychiatric disorders. (**A**) Volcano plot showing log_2_ Fold Change and −log_10_ adjusted p-value for variants that have enhancer activity from at least one allele. Significant variants (FDR < 0.1) were annotated with PLAC-seq predicted target gene name and color-coded based on target gene expression in mid-gestation telencephalon. Two vertical dashed lines indicate the absolute log_2_FC of 1. The horizontal dashed line indicates FDR at 10%. (**B**) Upset plot showing the number of variants (bar) passing combinations of different thresholds (dots and lines below bar). The number of DAVs was highlighted in red. (**C**) Enrichment log_2_ odds ratio of DAVs overlapping different features, including combined or separate cell-type-specific DA regions, adult brain eQTL, GWAS of various psychiatric disorders and low-frequency variants with minor allele frequency (MAF) less than 0.01. (**D**) TFBSs predicted to be altered by DAVs using motifbreakR. Dot color represents TF expression in primary cortical cells, size represents predicted magnitude of binding affinity alternation. TFs were ranked by TFBS alternation significance (-log_10_p-value, y-axis). (**E**-**F**) Genomic browser tracks showing examples of causal regulatory variants and their predicted target genes. The top track shows PLAC-seq chromatin loop in EN (*32*), the second track shows bulk RNA-seq in primary cortical cells, the third track shows bulk H3K27ac ChIP-seq (*31*), followed by a track of bulk ATAC-seq in deep-layer cortex (*31*). The bottom ten tracks show scATAC-seq in human cortex (*11*). DAV rs2193495 (**E**), located in a dlEN-specific accessible region, potentially down-regulates *TBR1* expression due to the introduction of EOMES and MAZ binding sites. DAV rs2154984 (**F**) is predicted to regulate *MARK2* expression and disrupt MTF1 and ZNF148 and introduce PPARD and NR2F6 binding sites. (**G**) Manhattan plot of SCZ-associated chromosome band 6p21.2 showing the 38 variants tested. The y-axis shows −log_10_ of adjusted p-value from MPRA. DAVs are highlighted in red and annotated with their predicted target gene. Arrows indicate the direction of allele effect observed in MPRA. (**H**) DAV rs9368977 located in 6p21.2 is predicted to disrupt binding of SP3, SP1, KLF4 and EGR4. (**I**) Manhattan plot of ASD-associated chromosome band 16p11.2 showing the 25 variants tested. The y-axis shows −log_10_ of adjusted p-value from MPRA. DAVs are highlighted in red and annotated with predicted target genes. The arrow indicates the direction of allele effect observed in MPRA. (**J**) TFBS altered in rs145650870. The alternative allele favors the binding of EHF and ELK3.

Next, we compared our DAVs to prior studies. A recent MPRA for dementia-associated variants in human embryonic kidney cells (HEK293T) (*51*) included 96 variants that were also in our library. We found that 89 variants show no significant allelic effect in either study, and 7 altered enhancer activity in HEK293T cells but not in our primary cortical cell data. This difference could be due to the cell types and/or the thresholds used to assign differential activity. Comparing our DAVs to eQTL data from psychENCODE (*34*), we found that 55% of DAVs (n = 77) had effects in the same direction as the eQTL and effect sizes were weakly correlated (Pearson’s correlation = 0.14, p = 0.102) (**fig. S5A**). Despite this, the correlation between MPRA and eQTL in non-DAVs (Pearson’s r = 0.008, p = 0.366) was notably lower than that in DAVs (**fig. S5A**). This corroborates that our lentiMPRA is efficient in identifying functional variants while underscoring differences between reporter activity and endogenous gene expression.

To decode the mechanisms through which the 164 DAVs exhibit differential activity, we predicted losses and gains of TFBS motifs using motifbreakR (*54*) (threshold = 1e-5), identifying 34 DAVs (21%) in which the alternative allele alters at least one TFBS (**Fig. 3D**). We also found that there is significantly more TFBS disruption compared with non-DAVs (OR = 1.49, p = 0.047, Fisher’s exact test). We then analyzed whether these predicted disrupted TFs functionally or physically interact with each other using the STRING database (*55*) and found a significant TF network centering on *SOX2* and *STAT3* (PPI enrichment p < 1e-16, **fig. S5C**).

We next predicted the putative target gene/s of DAVs using chromatin interaction data in various brain cell types (*32*, *33*, *56*) and adult brain eQTLs (*34*, *36*)(**Fig. 3A**), finding 48 DAVs (29.3%) to have chromatin loops with gene promoters and 8 of these (17%) to be eQTLs for the interacting gene. As regulatory activities vary over development, target genes predicted using adult brain eQTLs may not reflect genes regulated in early brain development, and thus we prioritized target genes predicted from chromatin interaction data. Many target genes are known risk genes or within susceptibility loci for psychiatric disorders and neural diseases. For example, we found that variant rs2193495, located in a dlEN-specific DA region that is thought to regulate the expression of T-box brain transcription factor 1 (*TBR1*), an ASD haploinsufficient associated gene and neurodevelopmental regulator (*57*), leads to reduced MPRA activity, possibly due to the creation of EOMES and MAZ binding sites (**Fig. 3E**). Another down-regulating variant, rs2154984, resides in a putative enhancer of the microtubule affinity regulating kinase 2 (*MARK2*), a risk gene whose loss-of-function variants have been associated with ASD (*58*) (**Fig. 3F**). This variant decreases the affinity of MTF1 and ZNF148 binding sites while increasing the affinity of PPARD and NR2F6 binding sites (**Fig. 3F**). Another example includes SCZ-associated variant rs10786689 that is thought to regulate nuclear factor kappa B subunit 2 (*NFKB2*) and suppressor of fused homolog (*SUFU*). This variant decreases enhancer activity, possibly due to the disruption of a SOX2 and/or SOX4 TFBS (**fig. S5D**). Since both genes were found to be up-regulated in SCZ patients (*59*, *60*), our results suggest that the alternative allele of rs10786689 could be protective. Finally, rs73392121 resides within a microglia scATAC peak and is thought to regulate NPC intracellular cholesterol transporter 1 (*NPC1*), a known cause of Niemann-Pick disease type C, with the alternative allele leading to reduced MPRA activity. Mutations in this gene lead to impaired cholesterol and lipid cellular transport, including microgliosis (*61*). These findings demonstrate that lentiMPRA can nominate candidate causal variants for known disease genes.

As a second strategy for linking DAVs to psychiatric disorders, we focused on known risk loci with multiple variants tested in our lentiMPRA. For example, in the SCZ-associated region 6p21.2 (*62*) (**Fig. 4G**), we tested 38 variants and found 2 DAVs: rs6912602 and rs9368977. DAV rs6912602 is one of the most differentially active in our lentiMPRA (3.3-fold decrease) and is an eQTL associated with reduced expression of peptidylprolyl isomerase like 1 (*PPIL1*). Partial loss-of-function variants in *PPIL1* cause neurodegenerative pontocerebellar hypoplasia in humans and mice (*63*). DAV rs9368977 increases activity in lentiMPRA, resides in open chromatin in RG and IPC, and is an eQTL for chromosome 6 open reading frame 89 (*C6orf89*). The alternative allele of rs9368977 disrupts the motifs of USFs SP3 and KLF4 (**Fig. 4H**). In the SCZ-associated locus 6p21.1 (*64*), we tested 48 variants and identified one DAV: rs1343025. The alternative allele of rs1343025 is associated with increased expression of vascular endothelial growth factor A (*VEGFA*). *VEGFA* regulates cerebral blood volume and is associated with SCZ, though the exact impact of *VEGFA* remains controversial (*65*). Another example, is the ASD risk loci is 16p11.2 (*66*), where we tested 25 variants and discovered one activity-increasing DAV, rs145650870 (**Fig. 4I**). This variant is located in an RG-specific chromatin loop for three nearby genes: tu translation elongation factor (*TUFM*), ataxin type 2 (*ATXN2L*) and SH2B adaptor protein 1 (*SH2B1*). The alternative allele of rs145650870 creates a TFBS for EHF (**Fig. 4J**). Combined, these results show that our lentiMPRA approach could be used to prioritize variants that have an effect on regulatory activity in disease-associated loci.

**Fig. 4.**
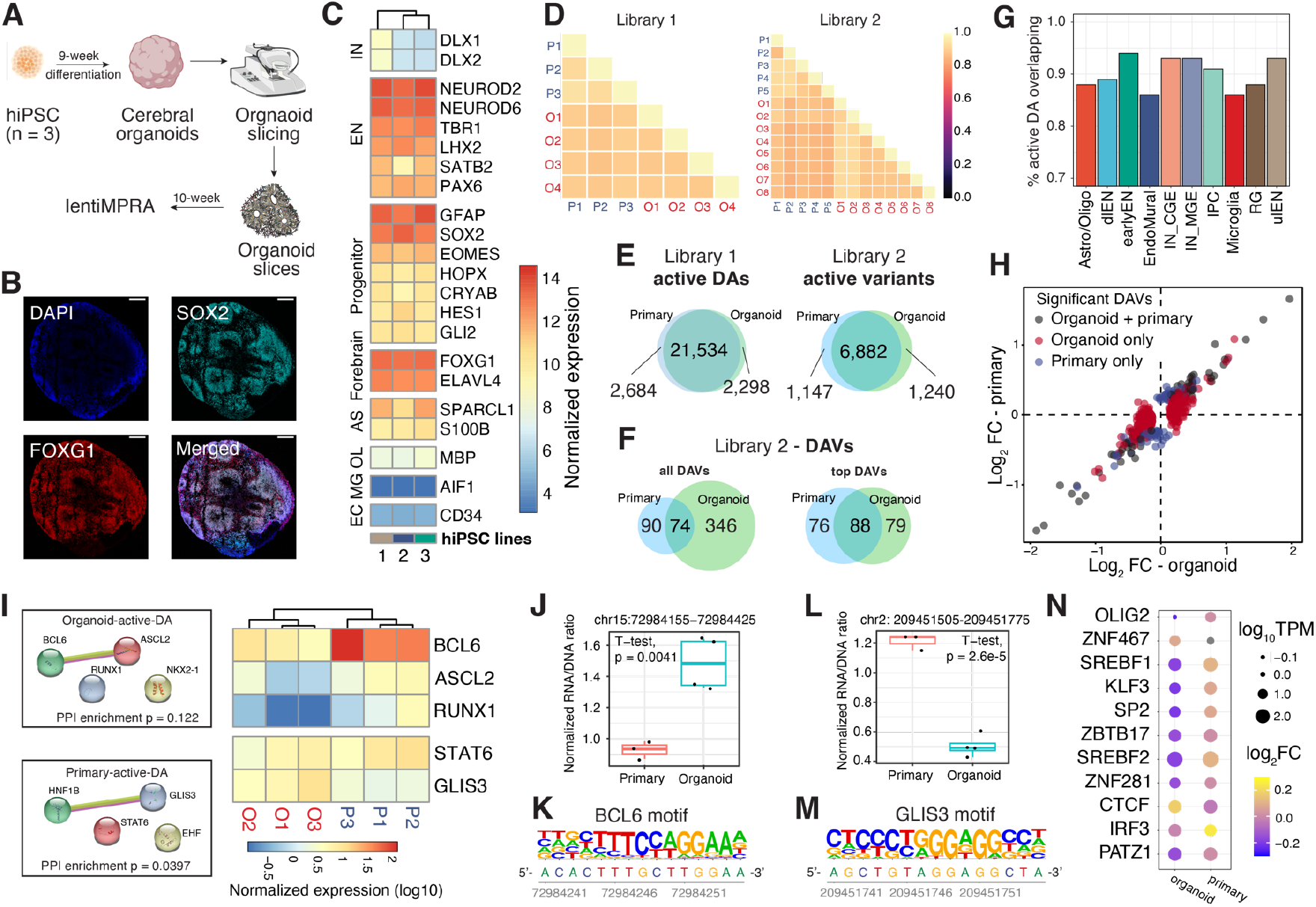
Comparison of lentiMPRA results in cerebral organoids and primary cortical cells. (**A**) Schematic of the experimental workflow. (**B**). Microscopic images of 10-week-old organoid slices immunostained for SOX2 (Cyan), FOXG1 (Red), and DAPI. Scale bar, 200 μm. (**C**) Normalized transcript count of marker genes in organoids derived from 3 hiPSC lines (1 = 21792A, 2 = 1323_4, 3 = 20961B). (**D**) Correlation of log2(RNA/DNA) between replicates in organoids and primary cortical cells for library 1 and 2, respectively. (**E**-**F**) Venn diagrams showing the overlap between organoids and primary cells. (**E**) Left: overlap of active DA regions; Right: overlap of active variants. (**F**) Overlap of DAVs. ‘Top DAVs’ were identified using shuffled sequences to define active and applying a cutoff for absolute log_2_FC of 0.3. (**G**) The proportion of active DAs in organoids that are also active in primary cells. (**H**) lentiMPRA log_2_FC in organoid (x-axis) and primary cells (y-axis). The scatter plot includes variants identified as DAVs in both organoids and primary cells (grey), variants detected as DAVs only in organoids (red), and variants detected as DAV only in primary cells (blue). (**I**) Left: Protein-protein interactions (PPI) of enriched TFBS motifs in active DAs specific to organoids or primary cells. PPI network generated using STRING (*73*) database. Right: heatmap showing the normalized transcript count of enriched TFs from bulk RNA-seq data. TFs not expressed (TPM < 1) in all replicates were removed from the heatmap. (**J-K**) A DAV (chr15:72984155-72984425, hg38) that contains a BCL6 motif showed increased activity in organoids versus primary cells (**J**) and its reference sequence contains a BCL6 motif (**K**). (**L-M**) A DAV (chr2: 209451505-209451775, hg38) with GLIS3 binding motif showing increased MPRA activity in primary cells versus organoids (**L**) and the location of the GLIS3 motif in its reference sequence shown below (**M**). (**N**) TFBSs altered by DAVs that show an opposite direction of allelic effect between organoids and primary cells. Dot sizes represent normalized TF expression; color represents log_2_FC.

### Organoids show comparable lentiMPRA activity to primary cells

Previous single-cell transcriptomic and epigenomic data along with immunohistochemical analyses suggest that cortical organoids recapitulate many of the cell types in the developing human forebrain (*11*, *46*, *67*–*69*). To explore the suitability of organoids as a tissue source for MPRAs, we tested both our lentiMPRA libraries in 10-week-old cortical organoids (**Fig. 4A**), which were validated for the expression of relevant cell type markers, such as *FOXG1*, *PAX6*, *EOMES*, *LHX2*, via immunostaining (**Fig. 4B**) and bulk RNA-seq (**Fig. 4C**). Following nine weeks of directed differentiation towards a dorsal forebrain fate, organoids were sectioned into 300-μm-thick slices and infected with the lentiMPRA libraries at 10 weeks. This slicing approach allowed diffusion of lentivirus into the majority of cells, providing high integration rates per cell (Multiplicity of infection (MOI) = 100). Slicing is also known to attenuate hypoxia, leading to better organoid cell health (*70*). For each library, we infected organoids derived from 2-3 iPSC lines with 2-4 technical replicates each and analyzed the data as described above for primary cells. We observed a high correlation between lentiMPRA replicates (average Pearson correlation over all tested sequences in each library, DA: 0.89, variant: 0.90) and positive controls consistently showed higher enhancer activity compared to negative controls (DA: p = 6.6e-4; variant: p = 0.027, Wilcoxon test, **fig. S2A**), confirming the high quality of our organoid data.

We compared RNA/DNA ratios between organoids and primary cortical cells and observed high correlation for both libraries (average Pearson correlation DA: 0.89, variant: 0.87, **Fig. 4D**). Similar to primary cells, roughly half of tested sequences were active (total: 31,954, DA: 23,832, variant: 8,122). The vast majority of organoid active sequences were also active in primary cells (**Fig. 4E**). To put this high level of concordance in the context of gene regulation, we performed bulk RNA-seq on three primary and three organoid samples (average replicate Pearson correlation, primary: 0.98, organoid: 0.99) and observed similar transcript levels between primary and organoid samples (average Pearson correlation 0.88), with some notable exceptions that we discuss below. Finally, we compared the activity of DA sequences stratified by the cell types in which they are accessible and found that active DA sequences were highly concordant in organoids versus primary cells for each cell type (**Fig. 4G**). The two cell types in which the lowest proportion of primary cell active DAs replicated in organoids were microglia (86.1%) and endothelial cells (86.4%), which is expected since these cell types are thought to be absent in cerebral organoids and our ability to assay activity of these DAs relies upon the permissiveness of MPRAs. These results suggest that cerebral organoids are a reasonable *in vitro* model of mid-gestation enhancer activity and gene expression, despite some differences in cell type composition.

Next, we examined the concordance of differential allelic activity between organoids and primary cells. In organoids, we observed a median absolute log_2_AR of 0.066, similar to that in primary cells (0.069), and detected 420 DAVs (FDR<10%), of which 74 (18% of organoid DAVs, 45% of primary DAVs) were also DAVs in primary cells (**Fig. 4F**). The larger number of DAVs identified in organoids is due to additional replicates and smaller batch effects. Consistent with this, the overlap of DAV sets was higher (53% of organoid DAVs, 54% of primary DAVs) when considering only the most differentially active organoid variants (absolute log_2_AR > 0.3 and activity above the median of shuffled controls, **Fig. 4F**). Despite this modest concordance in which variants were statistically significant, we observed a high correlation in DAV effect sizes in organoids versus primary cells (r = 0.91, p = 2.2e-16, **Fig. 4H**). We conclude that cerebral organoids and primary cells produce comparable lentiMPRA measurements of differential allelic activity for variants with the largest effects, with noise and cell type differences affecting measurements at and below the significance threshold for identifying DAVs.

Beyond evaluating the suitability of organoids for modeling primary cells, we were also curious about what differences in lentiMPRA results between the two settings would reveal about the biology of early neurodevelopment. Focusing first on the 2,298 DA sequences that were active only in organoids and 2,684 only in primary cells, we performed motif enrichment analysis and examined the expression level of enriched TFs (**Fig. 4I**). Organoid-specific active DA sequences were enriched for binding sites of NKX2.1, RUNX, BCL6, and ASCL2. *BCL6* is a transcriptional repressor with significantly lower expression in organoids compared to primary cells (q-value = 8.86e-7), consistent with our observation that sequences harboring BCL6 motifs tend to have higher lentiMPRA activity in organoids. One such example, includes a dlEN-specific accessible region containing a BCL6 motif that had significantly higher enhancer activity in organoids (**Fig. 4J-K**). In addition, overexpression of *BCL6* is known to inhibit apoptosis (*71*) and therefore could reflect elevated cell stress in organoids (*72*). For the primary-specific active DA peaks, we observed enrichment for GLIS3, STAT6, EHF, and HNF1B motifs. Compared to primary cells, organoids showed higher *GLIS3* expression (q-value = 9.24e-7) and we observed higher lentiMPRA activity in primary cells versus organoids for an Astro/Oligo and IN-MGE DA region containing a GLIS3 motif, suggesting that it may be functioning as a repressor in these primary-specific active sequences (**Fig. 4L-M**). Thus, motif analysis helped us identify TFs whose differential expression between primary cells and organoids is associated with significant shifts in enhancer activity, suggesting repressor versus activator roles for these TFs and underscoring their importance in regulating neurodevelopment.

Next, we examined variants identified as DAVs only in organoids or primary cells. As most variants still showed highly comparable effect sizes in both, we focused on 61 variants showing an opposite direction of effect. We predicted TFBS losses and gains using motifbreakR and compared TF expression using bulk RNA-seq (**Fig. 4N**). Twenty-eight variants were predicted to alter TFBS and 50% of altered TFs showed differential expression between organoid and primary. For example, we found that variant rs112049982 increased enhancer activity in primary cells but decreased activity in organoids. rs112049982 was predicted to improve the binding affinity of OLIG2, a maker gene for oligodendrocytes (Oligo) and oligodendrocyte precursor cells (OPC), which showed significantly lower expression in organoids. This agrees with prior knowledge that Oligo and OPC are extremely rare populations in cerebral organoids (*11*). Thus, we speculate that the discordant effect of the alternative allele of rs112049982 in organoids versus primary cells could be due to *OLIG2’*s differential expression. Together, these results indicate that despite organoids being a suitable *in vitro* model, differences in the *trans*-regulating environment should be carefully examined when interpreting lentiMPRA results.

### A sequence-based deep learning model of lentiMPRA activity reveals neurodevelopmental enhancer motif grammar

Our large dataset of lentiMPRA measurements provides an opportunity to characterize the mid-gestation enhancer code by modeling enhancer activity and then decoding the model’s understanding of how sequence variants modulate activity. To do this, we first designed a deep learning regression model that combines a single convolutional layer to learn motif-like sequence features, followed by two recurrent layers to learn the position, spacing, and orientation of motifs (Methods). Sequences were one-hot encoded into matrices (270bp x 4 nucleotides per sequence), and the mean RNA/DNA ratio across replicates was used as the regression target variable. For each library, we trained a model on sequences from all chromosomes except chromosome 3 (used as a validation set to prevent overfitting during training) and chromosome 4 (held out completely for an independent measure of predictive performance). Controls were included in model training. The variant library also included 15,000 sequences that represent a range of expected activity levels due to varying epigenetic similarity to validated brain enhancers in the VISTA database (*74*). On chromosome 4, the DA and variant models achieved 0.82 and 0.78 Pearson correlation, respectively (**Fig. 5A**; 0.81 and 0.7 Spearman correlation). Other similar sequence-to-activity models include MPRA-DragoNN (*75*) trained on human HepG2 and K562 Sharpr-MPRA data (0.28 Spearman correlation), and DeepSTARR (*18*) trained on fruit fly STARR-seq data (0.68 Pearson correlation). Though direct comparisons are not possible due to vast differences in assay type and dataset quality, our held-out predictive performance suggests our model is learning relevant sequence features for predicting MPRA activity.

**Fig. 5.**
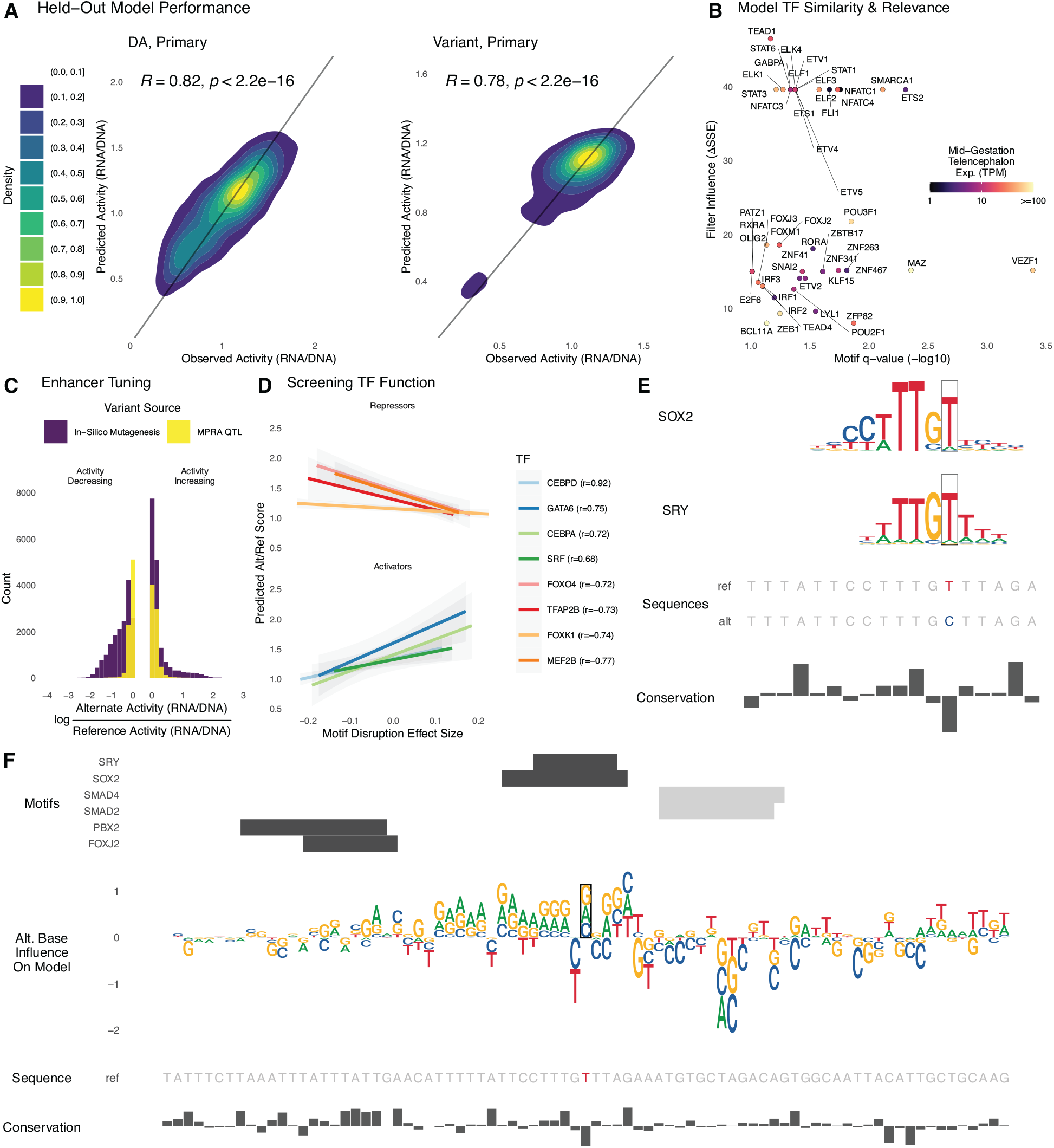
Sequence determinants of lentiMPRA activity can be modeled with deep learning. For each library, we trained a deep learning model to predict lentiMPRA activity in primary cells from sequence alone. (**A**) Sequences on chromosome 4 were held out from model training and used to evaluate model performance. Predicted and measured activity have high Pearson correlation for the DA library (left) and variant library (right). (**B**) The model learned motifs of neurodevelopmental TFs and used them for accurate predictions. Predictive importance of convolutional filters (change in sum of squared errors when fixing filter output to zero) is plotted against significance of matches to HOCOMOCO motifs (TOMTOM q-value < 0.1) for TFs expressed in mid-gestation telencephalon (mean CPM > 1). (**C**) Applying ISM to the variant library, we found that the activity of most enhancers can be tuned up and down through introduction of alternative alleles. The largest activity-increasing and activity-decreasing alleles for each sequence (purple) tend to have bigger effects than the lentiMPRA measured effects for QTLs (yellow). (**D**) We combined ISM with motifbreakR TFBS disruption scores to screen TFs for repressor versus activator function in neurodevelopment, using the most activity-increasing alternative allele for each sequence in the variant library. TF’s where predicted activity is anti-correlated with motif score tend to repress expression (top) and those with a positive correlation tend to be known activators (bottom). This relationship can be used to decode if the model has learned an activator versus repressor role for TFs that function in both ways. (**E**) The reference T allele of eQTL rs2883420 (lentiMPRA RNA/DNA 0.8) matches motifs of repressors SRY and SOX2, while the alternate C allele disrupts a high information content position in both motifs, resulting in a large activity increase (lentiMPRA RNA/DNA = 0.97, predicted RNA/DNA = 0.96). (**F**) ISM predicts that the other two possible alleles at rs2883420 also increase activity (middle, sequence logo indicates magnitude and direction: up=increasing, down=decreasing). Alternative alleles at adjacent nucleotides overlapping TF motifs (top, positive strand = black, negative strand = gray) have even larger predicted effects on activity. Region shown is chr10:86,851,230-86,851,500 (hg38).

Convolutional neural networks learn *de novo* filters from DNA sequences that represent position-specific nucleotide frequencies, similar to TFBS motifs. We therefore used the set of sequences that strongly activate each filter to construct a position weight matrix (PWM) and compared these against the HOCOMOCO (v11) database (*76*) to identify significant matches to know binding site motifs (Methods). As many filters have significant matches to motifs (q-value < 0.1) for TFs that are expressed in mid-gestation telencephalon (mean TPM > 1), we estimated each filter’s importance for predicting lentiMPRA activity by setting its output to zero and quantifying how much model performance decreases (deltaSSE; Methods). Top-ranked filters that match TFs with high mid-gestation telencephalon expression included TEAD1, NFATC1, STAT3, FOXJ3, POU2F1, and BCL11A (**Fig. 5B**). In addition to these TFs, our method also highlighted several USFs that function as cofactors to improve chromatin accessibility (*47*), consistent with our finding that motifs for these TFs are enriched in active versus inactive sequences (**Fig. 1D**). Thus, our model learned that both universal co-factors and TFs specific to the physiological context of our lentiMPRA experiments contribute to enhancer activity.

To complement this analysis, we performed a large-scale *in silico* mutagenesis (ISM) study. This method enabled us to quantify how individual nucleotide variants affect model predictions and did not directly rely upon a PWM database, though we did use PWM similarity to interpret high-scoring variants. Specifically, we constructed sequences with each possible alternate base at each of the 270 positions in each of the 17,069 variant-containing oligos for a total of 18.4 million alleles. We then predicted the activity of each alternate allele. Examining the distribution of the largest predicted activity change (up or down) per oligo, we found that while the QTLs tested in our lentiMPRA generally have moderate effects on activity, many of the adjacent synthetic variants tested with ISM have larger effects (**Fig. 5C**). For 11.6% of oligos, predicted activity can be increased by 50% or more through a single nucleotide change. Conversely, 19.7% of oligos can be reduced by 50% or more through a single change. As expected, activity-increasing variants frequently create binding sites for transcriptional activators (e.g., CEBPD) or mutate binding sites for repressors (e.g., FOXK1) that are expressed at mid-gestation, while activity-decreasing variants do the opposite (**Fig. 5D-E**). All sequences contained both increasing and decreasing alleles, and in most cases the two variants with the largest absolute ISM scores have opposing effects on activity. At nucleotides with large absolute ISM scores, the three alternative alleles tend to all be increasing or decreasing as expected if the reference base is a high information content position in a TFBS (**Fig. 5F**).

As an example, we highlight the region around eQTL rs2883420 (**Fig. 5E-F**) that has strong matches for SRY-like motifs. ISM predicts that all three alternative alleles at rs2883420 increase activity (predicted RNA/DNA ~ 0.97). In lentiMPRA, the reference allele was inactive in lentiMPRA (RNA/DNA ~ 0.8), while the alternative allele made the sequence nearly active (RNA/DNA ~ 0.96), fitting with our prediction. Further examination of the sequence effects of this eQTL (**Fig. 5F**), found a strong disruption of motifs for repression-capable TFs, such as SOX2 (*77*) and SRY (*78*) (p-value < 1e-4). ISM also predicts increased activity for non-reference alleles in a TFBS-sized region surrounding rs2883420, and most of these have larger effects than the eQTL, consistent with our genome-wide observations (**Fig. 5C**). These findings indicate that our model is learning *de novo* PWM-like representations of TF motifs which together form a neurodevelopmental regulatory grammar. Such a model can be leveraged to perform ISM, revealing how variants not present in an MPRA library alter enhancer activity and transcription factor binding. This strategy could be extended to discover and design cell-type specific enhancers with precisely tuned activity levels.

## Discussion

Gene regulatory elements have a major impact on human brain development and neurodevelopmental disease. Here, we combined lentiMPRA and deep learning to annotate thousands of regulatory elements in the developing cortex and cerebral organoids. This work provides a large catalog of functional human brain developmental enhancers and disease-associated variants, along with deep learning models that can accurately predict cell-type-specific regulatory regions and variant effects. In addition, it showcases the usability of cerebral organoids as an *in vitro* model for testing regulatory activity in mid-gestation, but also highlights several differences in the *trans*-regulating environment that should be taken into account when interpreting these results.

One caveat of our lentiMPRA is that it is a bulk assay with limited capability to detect cell-type specific signals. For example, we found that abundant cell types, such as neurons and radial glia, had higher percentages of active cell-type specific DA sequences compared to rarer cell populations. For microglia in particular, this could also be due to it being a difficult cell type to infect with lentivirus (*79*), leading to its lower active DA percentage (43.9%). Our validation of eleven regions for cell-type-specific activities in developing brain tissues identified three excitatory neuron-specific enhancers showing expected cell-type specificity, while the rest were active but non-specific. Conversely, we observed lentiMPRA activity for some microglia- and endothelial-specific DA sequences in organoids, despite these cell types being absent or very rare (*11*). We hypothesize this is due to sequences being activated by TFs present in other cell types that do not activate the endogenous sequence due to repressive chromatin. Beyond enhancer assays being permissive and testing sequences outside their endogenous context, other factors that may contribute to unexpected findings for sequences with cell-type specific chromatin accessibility include the limited resolution of scATAC-seq and differences in developmental stages. Nonetheless, nearly half of the 48,861 cell-type specific open chromatin regions that we tested were active enhancers in primary cells and/or organoids.

The cerebral organoids we generated from three hiPSC lines produced lentiMPRA measurements of enhancer activity that were highly correlated with measurements for the same sequences in primary cortical cells. While differential allelic activity was highly correlated for the variants with the largest effects, at least half of the DAVs identified in organoids or primary cells were not statistically significant in the other context with some having opposite allelic effects. We mostly attribute this to differences in sample sizes and cell type proportions. However, we also found that discordant results between organoids and primary cells could shed light on differences in the cellular environment between these two contexts. We identified *BCL6* and *GLIS3* as TFs whose differential expression in primary cells versus organoids can explain why sequences containing their TFBS motifs show significantly differential activity in lentiMPRAs. By looking at whether motifs are positively or negatively correlated with activity, both this analysis and our deep learning-based ISM analysis showed how lentiMPRA data can be used to infer if TFs are acting as repressors versus activators. These computational inferences are needed, because many TFs have both repressive and activating functions (e.g.(*80*–*82*)), and neurodevelopmental enhancers can play both activating and repressing roles depending on the bound TFs (*46*).

Another clear signal in our computational analyses was the importance of USFs for enhancer activity in early neurodevelopment. Stripe TFs are known to function as transcriptional co-factors improving chromatin accessibility (*47*). Motif enrichment in our lentiMPRA active sequences identified universal stripe factors alongside many known neurodevelopmental TFs. Corroborating this finding, analysis of our machine learning models identified many predictive features matching stripe motifs. These results suggest that stripe TF motifs may be useful for boosting the activity of designed enhancers, while they might reduce or overshadow signals from cell-type specific TFs. Investigations that study the role of universal stripe factors in MPRAs and *in vivo* will be an important direction for future studies.

We evaluated the regulatory effect of 17,069 brain QTLs and psychiatric disorder associated variants, identifying 164 differentially active variants. This small number is in line with other MPRAs that tested the effect of single-nucleotide variants (*14*, *51*, *53*, *83*, *84*) observing relatively small effects of single nucleotide substitutions, especially common alleles, on regulatory activity. Our deep learning model supports this conclusion; it predicts that many nucleotide changes in the open chromatin regions we tested, including alleles never or rarely seen in people, would show greater differential activity than the brain QTLs did. Another contributing factor is that regulatory variants, unlike coding variants, impact different layers of transcriptional regulation. MRPAs detect variants affecting enhancer activity or perhaps TF binding (*85*) but not those that modulate chromatin states, genome folding, splicing, or other aspects of gene expression. Finally, since about half of the DAVs we detected are in cell-type specific open chromatin regions, we expect that performing lentiMPRA on mixed cell populations limits detection of allelic effects that vary across cellular contexts.

Despite detecting only 164 high-confidence DAVs, integrative analysis of our data with publicly available chromatin interaction data linked many of these DAVs to one or more target genes expressed in neurodevelopment. Predicted target genes of many DAVs are known risk genes or within susceptibility loci, such as *TBR1* and *MARK2* for ASD or *NFKB2* and *SUFU* for SCZ. In particular, for large psychiatric disorder associated loci, our results for 6p21.1, 6p21.2 and 16p11.2 showcase the utility of lentiMPRA to identify potential disorder-associated regulatory variants in a high-throughput manner. In summary, we nominated several differentially active QTLs as potential causal variants of known disease genes/loci, paving the way for developing novel genetic diagnostic and therapeutic tools.

Overall, our work strengthens the utility of using primary cell culture, organoids, and MPRAs to investigate regulatory elements and variants involved in human brain development. In future work, it would be interesting to utilize an organoid lentiMPRA approach to test libraries from various psychiatric disorder-derived iPSCs to identify donor-specific *trans* effects on regulatory activity. iPSC derived organoids are also a promising avenue for investigating brain development in non-human primates, including comparative studies. Another technological development that could be used to expand upon this study is single-cell MPRA (*86*, *87*). While currently limited to a small number of sequences, this approach could eventually overcome some limitations we faced testing cell-type specific DAs in a bulk assay. It will also be critical to leverage CRISPR screens to assess the endogenous activity of candidate enhancer sequences, including those validated for activity with MPRAs. While CRISPR-based tools will likely improve our understanding of regulatory element cell-type specificity, they have their own caveats such as the need for high effect sizes on target gene mRNA levels. Deciphering the regulatory code of human brain development will require integration of all these strategies. The datasets and models generated in this work are a step in that direction.

## Acknowledgments

**General**: We would like to thank members of the Ahituv, Nowakowski, Pollen and Pollard labs for assistance with this manuscript. We would like to thank Kevin White and Sophia Gaynor for sharing control sets of MPRA. Data were generated as part of the PsychENCODE Consortium, supported by: U01DA048279, U01MH103339, U01MH103340, U01MH103346, U01MH103365, U01MH103392, U01MH116438, U01MH116441, U01MH116442, U01MH116488, U01MH116489, U01MH116492, U01MH122590, U01MH122591, U01MH122592, U01MH122849, U01MH122678, U01MH122681, U01MH116487, U01MH122509, R01MH094714, R01MH105472, R01MH105898, R01MH109677, R01MH109715, R01MH110905, R01MH110920, R01MH110921, R01MH110926, R01MH110927, R01MH110928, R01MH111721, R01MH117291, R01MH117292, R01MH117293, R21MH102791, R21MH103877, R21MH105853, R21MH105881, R21MH109956, R56MH114899, R56MH114901, R56MH114911, R01MH125516, R01MH126459, R01MH129301, R01MH126393, R01MH121521, R01MH116529, R01MH129817, R01MH117406, and P50MH106934 awarded to: Alexej Abyzov, Nadav Ahituv, Schahram Akbarian, Kristin Brennand, Andrew Chess, Gregory Cooper, Gregory Crawford, Stella Dracheva, Peggy Farnham, Michael Gandal, Mark Gerstein, Daniel Geschwind, Fernando Goes, Joachim F. Hallmayer, Vahram Haroutunian, Thomas M. Hyde, Andrew Jaffe, Peng Jin, Manolis Kellis, Joel Kleinman, James A. Knowles, Arnold Kriegstein, Chunyu Liu, Christopher E. Mason, Keri Martinowich, Eran Mukamel, Richard Myers, Charles Nemeroff, Mette Peters, Dalila Pinto, Katherine Pollard, Kerry Ressler, Panos Roussos, Stephan Sanders, Nenad Sestan, Pamela Sklar, Michael P. Snyder, Matthew State, Jason Stein, Patrick Sullivan, Alexander E. Urban, Flora M. Vaccarino, Stephen Warren, Daniel Weinberger, Sherman Weissman, Zhiping Weng, Kevin White, A. Jeremy Willsey, Hyejung Won, and Peter Zandi. Sequencing was partially carried out by the DNA Technologies and Expression Analysis Core at the UC Davis Genome Center, supported by NIH Shared Instrumentation Grant 1S10OD010786-01.

## Author contributions

Conceptualization: CD, SW, PP, AP, TN, NA, KP

Methodology: CD, SW, RZ, PP, NA, KP, DA, DC, SN, FMV

Investigation: CD, SW, MS, RZ

Visualization: CD, SW, MS

Funding acquisition: NA, KP

Project administration: CD, SW, RZ, NA, KP

Supervision: TN, NA, KP

Writing - original draft: CD, SW, MS, NA, KP

Writing -review & editing: All authors

## Data and materials availability

The source data described in this manuscript are available via the PsychENCODE Knowledge Portal (https://psychencode.synapse.org/). The PsychENCODE Knowledge Portal is a platform for accessing data, analyses, and tools generated through grants funded by the National Institute of Mental Health (NIMH) PsychENCODE Consortium. Data is available for general research use according to the following requirements for data access and data attribution: (https://psychencode.synapse.org/DataAccess). For access to content described in this manuscript see: DOI will be provided prior to publication.

