## Supplementary Materials for "Massively parallel characterization of psychiatric disorder-associated and cell-type-specific regulatory elements in the developing human cortex"

Science
AAAS

Supplementary Materials for

### Massively parallel characterization of psychiatric disorder-associated and cell-type-specific regulatory elements in the developing human cortex

Chengyu Deng, Sean Whalen, Marilyn Steyert, Ryan Ziffra, Pawel F. Przytycki, Fumitaka Inoue, Daniela A. Pereira, Davide Capauto, Scott Norton, Flora M. Vaccarino, Alex Pollen, Tomasz J. Nowakowski, Nadav Ahituv, Katherine S. Pollard

**The PDF file includes:**

Materials and Methods
Supplementary Text
Figs. S1 to S5
Tables S1 to S2
References

**Other Supplementary Materials for this manuscript include the following:**

Data S1 to S3

#### Materials and Methods

##### lentiMPRA library design

Pseudo-bulked ATAC-seq peaks were computed (macs2 2.2.7.1, IDR 10%) on single cell ATAC-seq data from primary human brain samples (*11*) and used to design two libraries of candidate enhancer sequences. Candidate peaks overlapping ENCODE blacklist regions or promoters (*88*) were removed.

Library 1: We determined differentially accessible (DA) peaks per cell type (DiffBind 3.0). Candidate DA peaks were then required to overlap H3K27ac peaks from bulk prefrontal cortex (PFC) tissue (ChIP-seq, 15 gestational weeks (*31*)). Alternately, microglia and non-microglia DA peaks could overlap H3K27ac peaks from their respective cell types (CUT&Tag, (*11*)). Candidates lacking an H3K27ac peak overlap were still included if they overlapped a cell type-matched statistically significant chromatin loop in neurodevelopmental Promoter Capture Hi-C (*33*) or PLAC-seq (*32*) data lifted over to the hg38 human genome assembly. Finally, 270-bp oligonucleotides ("inserts") were created from ATAC-seq peaks by extending 135-bp in each direction from the peak summit.

Library 2: We first required that candidate ATAC-seq peaks (non-DA) overlap H3K27ac ChIP-seq peaks from bulk PFC tissue (15 gestational weeks) (*31*) or from at least one matched cell type in single-cell CUT&Tag data (*11*). Candidates were then required to be located within 100 kilobases of a differentially expressed cross-disorder neurodevelopmental gene (http://resource.psychencode.org) or overlap a quantitative trait locus (QTL) sharing a Linkage Disequilibrium (LD) block with a neuropsychiatric genome-wide association study (GWAS) SNP. Separate LD blocks were computed for each 1000 Genomes super-population (AFR = African, AMR = Admixed American, EAS = East Asian, EUR = European, SAS = South Asian) using bi-allelic SNPs with a minimum minor allele frequency of 5% (*89*). All QTL and GWAS coordinates were converted to hg38 by referencing their rsid in dbSNP (*90*). The reference allele was determined by examining whether allele 1 or 2 matched the reference hg38 genome sequence. Then, 270-bp inserts were created from ATAC-seq peaks by extending 135-bp in each direction from the overlapping QTL. Multiple inserts were created if multiple QTLs overlapped a peak. QTLs included GTEx high-confidence eQTLs from primary brain tissue (https://gtexportal.org), PsychENCODE eQTLs from primary brain tissue w/FPKM greater than 1 in > 20% of samples (*34*), PsychENCODE cQTLs from primary brain tissue (*34*), Gandal et al. eQTLs (*9*), Liang et al. cQTLs from primary neuron and progenitor cell lines (*35*), and Werling et al. eQTLs from primary brain tissue (*36*). GWAS SNPs were obtained from the Psychiatric Genomics Consortium for Attention Deficit Hyperactivity Disorder (*37*), Alzheimer's Disease (*38*), Autism Spectrum Disorder (*39*), Bipolar Disorder (*40*), Cross-Disorder (*8*), Major Depression Disorder (*41*), Obsessive-Compulsive Disorder (*42*), Schizophrenia (*43*), and Tourette Syndrome (*44*). Also included were inserts for Human Accelerated Regions (HARs, (*91*)), zooHARs (*92*), and peaks with low, medium, and high similarity to validated VISTA brain enhancers (*74*) based on epigenetic profiles.

Shared Controls: Both libraries included positive and negative controls from the Vaccarino lab and the White lab, as well as shuffled sequences (https://github.com/agordon/fasta_ushuffle) of inserts from their respective libraries.

**Primary cortical cell culture**

De-identified tissue samples were collected with previous patient consent in strict observance of the legal and institutional ethical regulations. Protocols were approved by the Human Gamete, Embryo, and Stem Cell Research Committee (institutional review board) at the University of California, San Francisco. Fresh cortical tissue was dissociated using papain (LK003150, Worthington Biochemical) into a single-cell suspension. After one DPBS wash and spinning at 300 rcf for 5 min, cells were then plated in PLO/laminin/fibronectin-coated 10cm dishes at a density of $3.5x10^5$ cells/cm2 and maintained in an incubator at 37°C, 5% CO2, and 8~10% O2. Culture medium (DMEM supplemented with 1x B27, 1x PSG, and 1x N2) was changed every 24 hours until lentivirus infection.

**Cortical organoid differentiation**

Prior to organoid differentiation, all iPSC lines were passaged in Stemflex (Gibco) + antibiotic-antimycotic (Gibco). Cortical forebrain organoids were generated as previously described (*11*, *72*, *93*). Briefly, confluent wildtype iPSCs (1323-4, 21792-A, 20961B) were dissociated using PBS+EDTA and split 1:3 into 6-well ultra low-attachment plates and maintained in a DMEM/F12-based induction media containing 15% KSR, 1% MEM-NEAA, 1% Glutamax, 1% Pen-Strep, 100 μM β-Mercaptoethanol, 5 μM Dorsomorphin, 5 μM SB-431542, 3 μM IWR1-endo, and 1X CEPT apoptosis inhibitor cocktail (50 nM chroman 1, 5 μM emricasan, 0.1% polyamine supplement, and 0.7 μM trans-ISRIB, per Yu et al, 2021(*94*)) for 6 days. Days 6-9, CEPT was not added. Aggregates were placed on an orbital shaker rotating at 80 rpm on day 9, then maintained in an expansion media containing a 1:1 mixture of DMEM/F12 and Neurobasal medium with 1% N2, 2% B27 without vitamin A, 1% Glutamax, 1% MEM-NEAA, 55 mM β-Mercaptoethanol, 1% Antibiotic-Antimycotic, 10 ng/mL FGF2, and 10 ng/mL EGF. EGF and FGF were not added starting on day 25. B27 containing vitamin A was used to replace the B27 without vitamin A starting on day 35. Media was supplemented with 10ng/mL LIF day 55-65, then 10ng/mL BDNF + 10ng/mL NT3 day 65 until fixation. Throughout culture, media was full-changed every 2-3 days. On day 60 post-aggregation, organoids were embedded in ACSF with 3% agarose solution then sliced at 300uM using a vibratome. Organoid slices were plated 10-to-an-insert on each millicell (Millipore) 6-well insert with 1mL media. After slicing, media was half-changed every 2-3 days, per Qian et al 2020(*70*).

**LentiMPRA**

<u>Construction and sequencing of plasmid libraries</u>

LentiMPRA was performed as previously described (*95*) with minor modifications. Briefly, an oligo pool containing both libraries was synthesized by Twist Bioscience. Each library was flanked by a unique 15bp adapter and thus was processed separately in the following steps. A 31bp minimal promoter and a 15bp random barcode were added to each 270bp candidate regulatory sequence through two rounds of PCR. The 485-bp amplicon was gel extracted and cloned by Gibson assembly (E2621S, NEB) into SbfI and AgeI-digested empty reporter backbone pLS-SceI (plasmid #137725, Addgene). Recombination products were transformed and amplified in electrocompetent cells (C3020, NEB) and grown in LB Agar plates (100217-214, VWR) at 37°C overnight. A 40,000 dilution was carried out in parallel to estimate the number of colonies in undiluted plates. Sixteen colonies were miniprepped (27106, Qiagen) from the dilution plate and Sanger sequenced to estimate the rate of error. >70% of colonies showed matched sequences without any mutations. We harvested ~3.5M colonies by Midiprep (12945, Qiagen). The number of colonies equals the number of oligos in the library multiplied by the desired number of barcodes for each oligo. 3.5M colonies yielded around 70 barcodes per oligo. After constructing the plasmid library, we generate a sequencing library to link each barcode to its corresponding oligo. A 477bp region in the plasmid containing the oligo and barcode was amplified using the following PCR conditions: combine 500ng of plasmid, 2µl of 100µM forward and reverse primers each, 100µl polymerase mix (M0544L, NEB), and $H_2O$ making up to 200µl, split into five PCR tubes and run the cycling program with an initial denaturation at 98℃ for 30s, 7 cycles of 98℃ for 10s, 65℃ for 75s, and a final extension at 65℃ for 5 minutes. The 540-bp amplicon was purified with x1 SPRIbeads (B23318, Beckman Coulter) and sequenced in one lane of an Illumina Nextseq Mid-Output PE150.

Lentivirus packaging, titration and infection

To package lentivirus, three million 293T cells were plated in a 15cm dish with 30 ml DMEM (10569044, ThermoFisher) with 10% heat-inactivated FBS (89510-194, VWR, heat-inactivated by incubating thawed serum at 56 ℃ for 30 min). The number of dishes needed depends on the library complexity and the infectability of the cells. Following two days of incubation, the plasmid library was transfected into 293T cells using lenti-pac kit (LT002, Genecopoeia) following the manufacturer’s protocol. Culture medium containing lentivirus was collected 48 hr post transfection. To concentrate lentivirus and achieve high titer, crude solution was centrifuged at 1,000g for 10 minutes to get rid of cell debris. Supernatant was then filtered through a 0.45 um PES unit (FB12566507, Fisher Scientific) to further remove dead cells. The flow-through was mixed with Lenti-X concentrator (631232, Takara Bio) in a 3:1 volume ratio and incubated at 4 ℃ for at least one day. Lentivirus was pelleted by centrifugation at 1,500g for 45 minutes at 4 ℃, and resuspended in ice-cold DPBS (D8537-6X500ML, Sigma Aldrich) with gentle pipetting. Concentrated virus was immediately stored at -80 in single-use aliquots.

To titrate lentivirus, 0.3-0.5 million primary cells were plated per well in a 24-well plate. Lentivirus was thawed at room temperature and used immediately after thawing. 0, 1, 2, 4, 8, 16, 32 and 64 µl of virus was added into each well separately. Fresh medium was replaced the next day. Genomic DNA was extracted (T3010L, NEB) from each well 72hr after infection. MOI was calculated by qPCR, as previously described (*95*). Titration on organoid slices were performed using 0, 10, 25, 75, 150 and 300 µl of virus per slice in 24-well plates.

For each replicate, 20 million primary cells dissociated from fresh cortex tissue at gestation week 18 were cultured in a 10cm dish for two days before infection. Based on titration results, each dish was infected with 300-500 µl of lentivirus library 1 or 2 to achieve an MOI of 85. Each barcode was estimated to be integrated into the host genome randomly around 250 times. Medium was refreshed the next day and cells were incubated for another two days before harvesting DNA and RNA. For organoids, 800 µl lentivirus library was added directly on top of the filter on day 70 of organoid slices at an estimated MOI of 100. Twenty organoid slices (roughly 7 million cells) were used per replicate. A full media change was performed on the next day. Organoids were cultured for another two days before being harvested.

DNA/RNA harvesting and sequencing

DNA and RNA were simultaneously extracted from infected samples using the Allprep kit (80204, Qiagen) following the manufacturer's protocol. Lysis buffer was prepared by adding 1% 2-mercaptoethanol to buffer RLT Plus. Briefly, primary cells in a 10cm dish were gently rinsed with DPBS once, then lysed in 1,200 µl lysis buffer and homogenized using QIAshredder columns (79656, Qiagen). Each organoid replicate was lysed with 2ml lysis buffer with vortexing until no cell clumps were visible. Purified RNA and DNA were quantified using Nanodrop. DNA was stored at -20℃ and RNA was stored at -80℃.

To prepare sequencing libraries, RNA was first treated with DNase to remove potential carry-over of gDNA using TURBO DNA-free kit (AM1907, Fisher Scientific). 7-9 µg RNA was reverse transcribed to generate cDNA using Superscript IV RT (Invitrogen; 18090200) and a custom designed RT primer (*95*). MPRA barcodes were amplified from 10-15 µg gDNA or cDNA separately through 2 rounds of PCR, which added the unique molecular identifier (UMI), index for demultiplexing and Illumina P5/P7 sequence. The following cycling program was used: initial denaturation at 98℃ for 2min, x cycles of 98℃ for 10s, 72℃ for 35s, and a final extension at 72℃ for 2min. The number of cycles needed is determined through a 10ul qPCR reaction. The ct value where fluorescence increase reached exponential phase was used as the number of cycles needed. Amplification reactions were cleaned up using SPRI beads at a ratio of x1.2. Library concentration was quantified via qPCR (NEB, E7630S). DNA and RNA barcode libraries were pooled in 1:3 mole ratio and sequenced with a NextSeq High-output SE75.

##### Culture of primary cortex slice for cell type specificity validation

Gestational week 16-22 tissue samples were initially maintained in ice cold artificial cerebrospinal fluid (125 mm NaCl, 2.5 mm KCl, 1 mm MgCl2, 2 mm CaCl2, 1.25 mm NaH2PO4, 25 mm NaHCO3, 25 mm d-(+)-glucose) bubbled with 95% O2 and 5% CO2. Samples were embedded in 3% low-melt agarose in ACSF, then sectioned to 300um thickness using a vibratome (Leica VT1200S). Each section included in this study contained full cortical thickness from VZ to CP. Lentiviral infections were performed by incubating slices in virus diluted 1:50 with media for 1 hour at 37°C. Slices were transferred to millicell (Millipore) inserts in a 6-well cell culture plate for maintenance at the air-liquid interface in 1mL of a media containing 60% Basal Medium Eagle (Gibco) 32% Hanks buffered saline solution (Lonza), 5% heat-inactivated FBS (Hyclone), 1% glucose (Sigma), 1% N2 (Thermo), 1% Glutamax (Gibco) and 1% Penicillin-Streptomycin (Gibco). Slices were maintained for 5 days at 37°C with 5% CO2 and 8% O2. Media was fully-changed 12 hours post infection, then half-changed every other day for the remainder of the five day culture period.

##### Immunocytochemistry and immunohistochemistry

Organoid and primary slice culture samples were fixed with 4% PFA in PBS for 1hr at 4C, then washed three times with PBS. Primary slices were stained floating in 200 µl of each solution, and washed with ~1mL PBS. Organoids were incubated in 30% sucrose overnight and embedded in a 1:1 mixture of OCT and 30% sucrose , then cryoblocks were stored at -80C until sectioning. Organoids were then cryosectioned to a 20um thickness. For staining, samples were blocked in a solution of PBS with 10% donkey serum + 0.2% gelatin + 0.1% TritonX for 30 minutes. Primary and secondary antibodies were diluted in blocking solution. Samples were incubated in primary antibody solution overnight at 4C, then washed three times with PBS at room temperature. Samples were then incubated in secondary antibody solution for 3hrs at room temperature, followed by a solution of PBS with 1ug/mL DAPI for 30 minutes and then washed 3 times with PBS before mounting samples with Fluoromount (Invitrogen). Primary antibodies in this study include: chicken anti GFP (1:1000, GFP-1020, Aves), mouse anti SATB2 (1:200, sc-518006, Santa Cruz), rabbit anti SOX9 (1:200, ab185230, Abcam), goat anti SOX2 (1:500, sc17320, Santa Cruz), rabbit anti HOPX (1:200, Santa Cruz). Secondary antibodies in this study include: donkey anti chicken 488 (1:500, A78948, Thermo), donkey anti mouse 647 (1:500, A31571, Thermo), donkey anti goat 647 (1:500, A32849, Thermo), donkey anti rabbit 594 (1:500, A11012, Thermo). All images were collected using 10x and 20x air objectives on a Leica SP8 confocal system, and processed using ImageJ/Fiji.

**LentiMPRA computational analyses**

Library quality control

Paired-end reads were merged using NGmerge (*96*) with a minimum overlap of 22 bp and maximum quality scores were used for error correction, for barcode associations as well as RNA and DNA. Association reads were aligned to their respective libraries using bwa mem 0.7.17 (*97*). DA library sequences required mapping quality > 30, while variant library reads set no mapping quality threshold due to the presence of alleles differing by one base. Sequences in both libraries required a CIGAR string of 270M. All barcode bases were required to have a minimum quality score of 30. Homopolymers and near-homopolymers were removed by requiring barcodes to have a Shannon entropy rate > 0.5. Barcodes needed to be observed at least three times, and to map to the same insert 90% of the time. Associations not matching these criteria were discarded. RNA and DNA barcodes and UMIs were required to be 15bp and 16bp, respectively. Barcodes were counted once per unique UMI to remove PCR duplicates.

Quantification

Inserts were required to have at least 10 unique barcodes (median DA primary: 56, DA organoid: 55, variant primary: 64, variant organoid: 64) and at least 40 total DNA barcodes. RNA and DNA were normalized for sequencing depth by converting raw counts to Counts Per Million (CPM), and activity for each insert was computed as RNA CPM / DNA CPM. Non-variant analyses used the mean RNA/DNA ratio across replicates; variants were analyzed using the activity of the alt allele divided by the reference allele, e.g. (alt RNA/DNA) / (ref RNA/DNA).

Batch correction

Variant library ratios were corrected to account for differences between samples from different donors (primary tissue) and batches (organoids) using limma 3.54 (*98*). Batch-corrected data was used in all analyses except testing for differential allelic effects (see below) where batch effects were instead accounted for by including a batch term in the design matrix. DA library ratios were from the same donor and batch and thus were not corrected.

Differential analysis

Differential expression between alternate and reference alleles was performed using limma 3.54 (*98*) on non-batch corrected ratios, including a batch term in the design matrix to avoid mis-estimating degrees of freedom. This resulted in 74 and 58 activity-increasing and decreasing variants at 1% FDR with absolute fold change > 1, with 87 increasing and 76 decreasing variants at 10% FDR with absolute fold change > 1.

##### Motif Enrichment

Motif enrichment was performed with HOMER 4.11.1 (*99*) using the median of positive controls to divide sequences into foreground (above) and background (below) sets, and the -h flag to use a hypergeometric test for p-values.

##### Gene Ontology Enrichment

GO enrichment was performed with g:Profiler (*100*) using the median of positive controls to divide sequences into foreground (above) and background (below) sets, and default parameters including a statistical test that accounts for the non-independent nature of terms in the GO hierarchy. Each insert in a set was mapped to the closest protein coding gene using bedtools (*101*) and biomaRt (*102*).

##### lentiMPRA Activity Modeling

A deep learning regression model was implemented in Tensorflow/Keras 2.11 (*103*) to predict lentiMPRA activity from sequence. The model was trained using the one-hot encoded 270bp DNA sequence of each oligo as features and the mean log RNA/DNA ratio as labels. A convolutional layer (relu activation, valid padding) first learns short motif-like features from DNA sequence, followed by a max-pooling layer to reduce dimensionality, followed by two Long Short-Term Memory layers to learn patterns of spacing and orientation; their outputs are flattened and connected to a dense output layer (linear activation). Inserts from chromosome 6 were held out as a validation set used for early stopping during training (patience 10), and inserts from chromosome 3 were held out as a test set for final performance evaluation (Spearman correlation). Inserts on all other chromosomes were used for 100 epochs of training with the adam optimizer and mean squared error loss. For each convolutional filter, importance was estimated by computing the sum of squared errors (SSE) for predictions made while fixing the filter's output to zero, and subtracting the SSE of the original predictions. Large increases in error indicate filters that are important to the model's accuracy.

##### In silico mutagenesis

In-Silico Mutagenesis (ISM) was performed on the variant library, creating alleles for all bases at each position for every sequence and using the model to compute an alt:ref ratio (AR). For each reference sequence, the allele with the largest activity-increasing (AR>1) and activity-decreasing (AR<1) predicted effects was tracked for each sequence. Using motifbreakR(*54*), we computed TFBS motif disruptions for the most activity-increasing allele per sequence and compared these to the predicted AR values. When alleles had opposing changes (AR decreases as motif score increases) we inferred that the model had learned a repressor role for the TF, while positive correlations suggest that the model learned an activator role. TFs with known, consistent roles as activators versus repressors were used as validation. Other TFs were screened for activator versus repressor activity in neurodevelopment.

##### Bulk RNA-seq analysis

RNA-seq was processed using Kallisto 0.48 (*104*) using default settings with Ensembl v96 annotations. Transcript abundances were aggregated into gene abundances using tximport 1.26.1 (*105*). Differential expression was computed using DESeq2 (*106*).

**Supplementary Text**

Predicted TFBS alternation showed a weak correlation with MPRA results

We observed a weak correlation between the predicted magnitude of TFBS alteration and lentiMPRA $log_2$AR ratio (Pearson's r = 0.16) in DAVs and a much lower correlation in non-DAVs (r = 0.0071). This low correlation is consistent with prior MPRA studies (*51*, *84*) and is driven in large part by the fact that only 44% of DAV-altered TFBS showed a consistent direction with lentiMPRA activity, perhaps due to the fact that these TFBS include motifs for both activators and repressors.

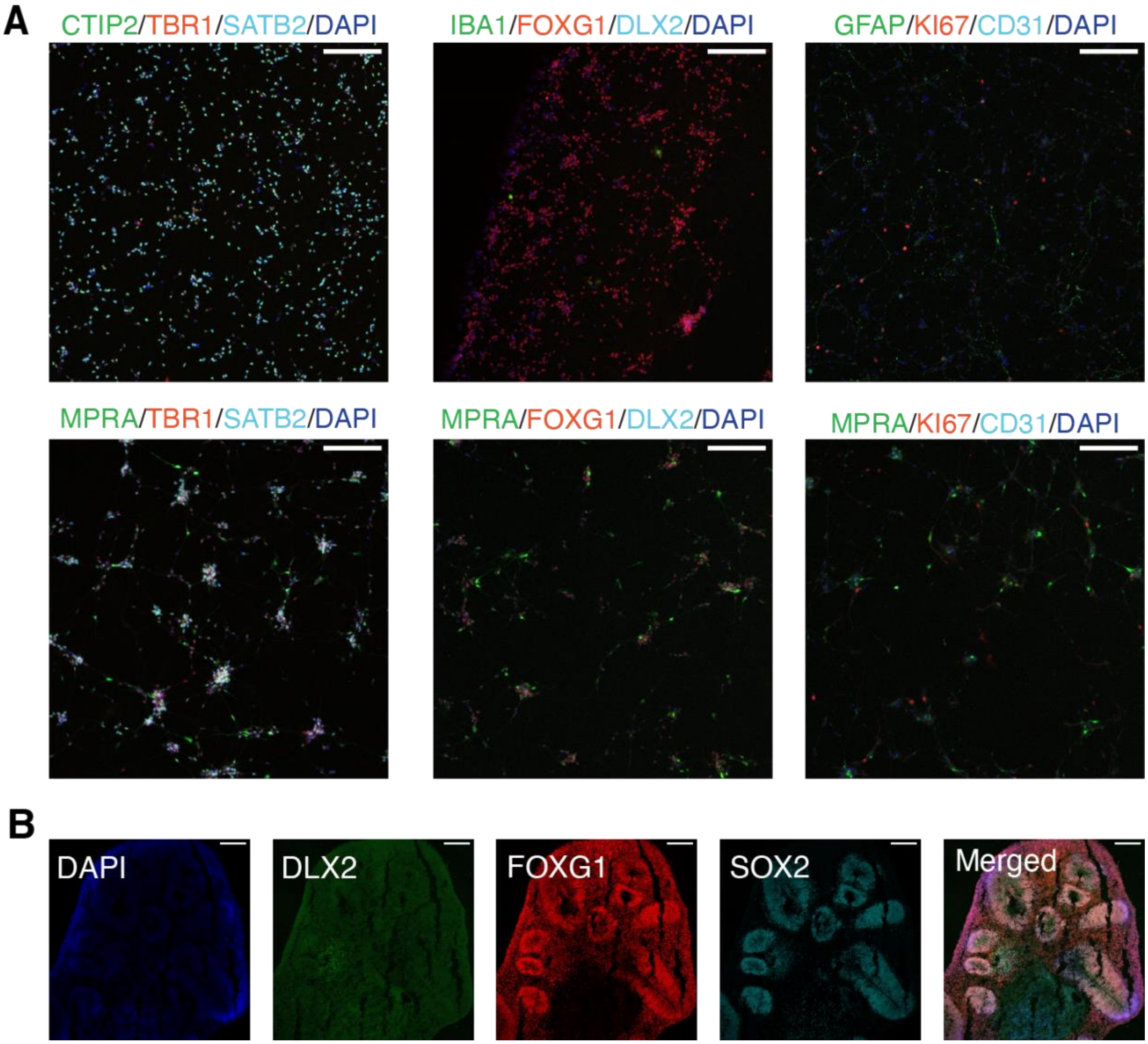


**Fig. S1. Primary cortical cell during mid-gestation and cerebral organoid characterization.** (**A**) Representative images showing primary cells dissociated from fresh human cortex at mid-gestation with immunocytochemical staining for various cell markers with (second row) or without lentiviral infection (first row). This confirms the presence of major cortical cell types and uniform infection across cell types. Scale bar, 200 µm. (**B**) Representative immunocytochemical staining images of 10-week-old cerebral organoids showing expression of progenitor and neuronal marker genes. Scale bar, 200 µm.

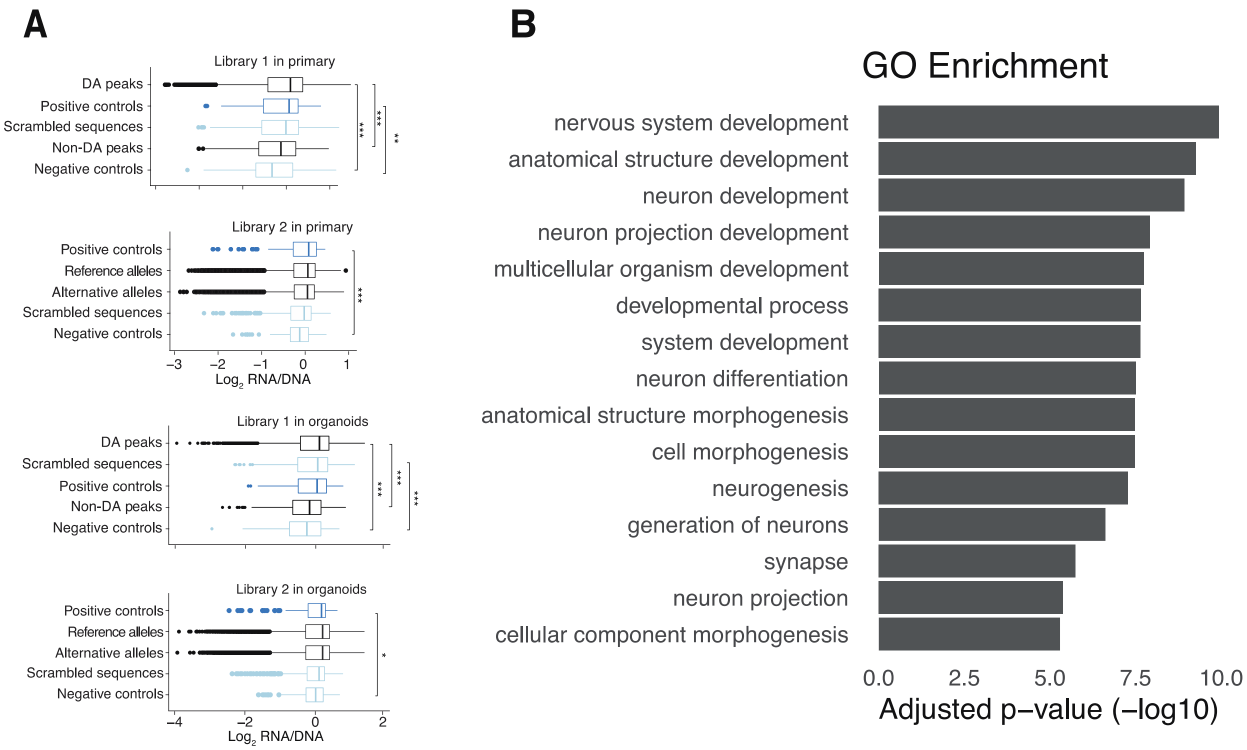


**Fig. S2. lentiMPRA distinguishes sequences with high and low regulatory activities.** (**A**) Boxplots showing the distribution of log2(RNA/DNA) in each sequence category. (**B**) Complete list of enriched GO terms from the 'Biological Process', 'Cellular Component' and 'Molecular Function' ontologies for nearest genes of the highest activity sequences (activity above the 75th percentile of positive controls; enrichment computed for both libraries combined). Closest genes of the lowest activity sequences (below 25th percentile) were used as a background set.

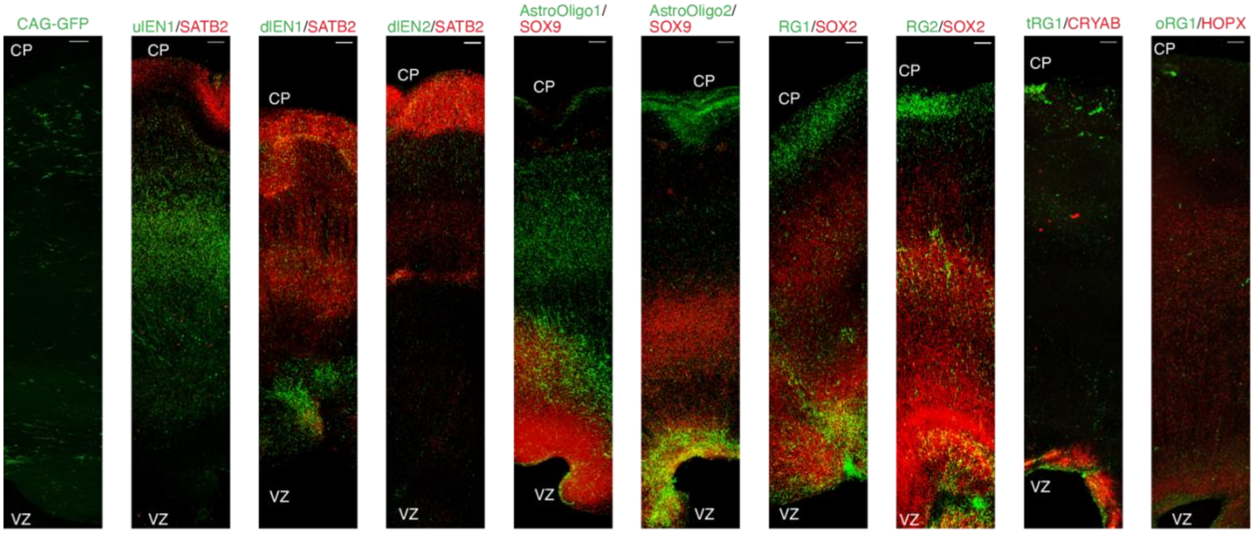


**Fig. S3. Validation of enhancers and their cell-type-specificity.** Mid-gestation human cortex slice cultures transduced with GFP lentivirus reporter driven by constitutively active enhancer CAG (left-most) or different candidate cell-type-specific enhancer (10x). Expression of GFP (green) and various cell markers (red) visualized via immunohistochemistry staining. Scale bar, 200 μm. The complete information is summarized in table S2. ulEN1, chr2:152408730-152409000; dlEN1, chr5:89274678-89274948; dlEN2, chr9:14297525-14297795; AstroOligo1, chr5:43217050-43217320; AstroOligo2, chr14:50985465-50985735; RG1, chr6:19663461-19663731; RG2, chr21:16154321-16154591; tRG1, chr1:68141016-68141286; oRG1, chr22:43331407-43331677 (all coordinates are human genome assembly *hg38*).

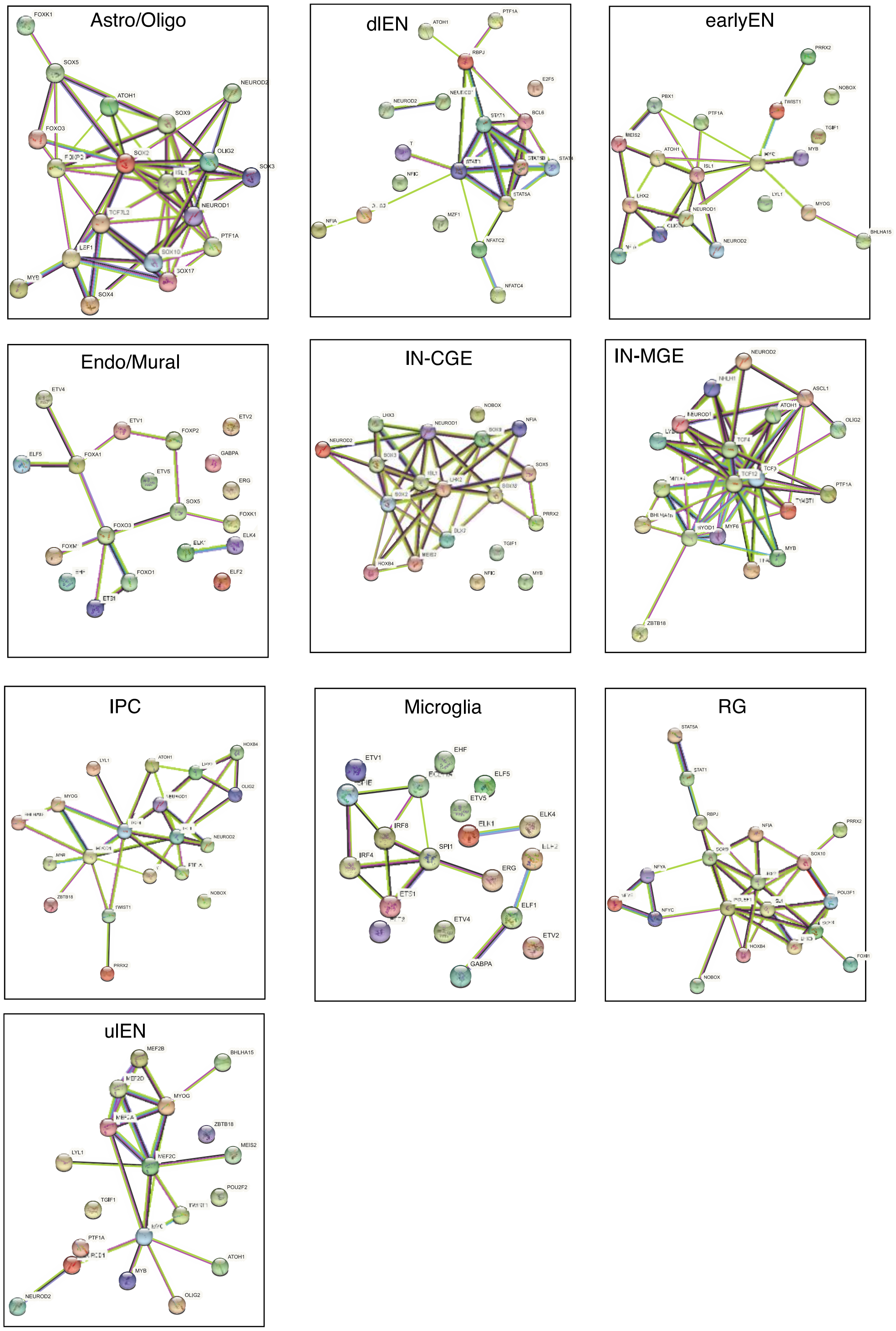


**Fig. S4. Active cell-type-specific DAs were enriched for TFs in significant protein-protein interaction (PPI) networks.** PPI networks from the STRING (*73*) database for TFs enriched in active DAs specific to different cell types. For each cell type, the top-enriched TFs in active DA regions tend to form significant functional networks, which could shed light on the mechanisms of cell fate determination during neurogenesis. For example, the top 20 enriched TFs of active DAs in IN-CGE formed a network centering on three genes in the *TCF* family, including *TCF3*, *TCF4* and *TCF12*. The *TCF* gene family, particularly *TCF4*, is known as a master transcriptional regulator during brain development (*107*), and a risk gene for SCZ (41).

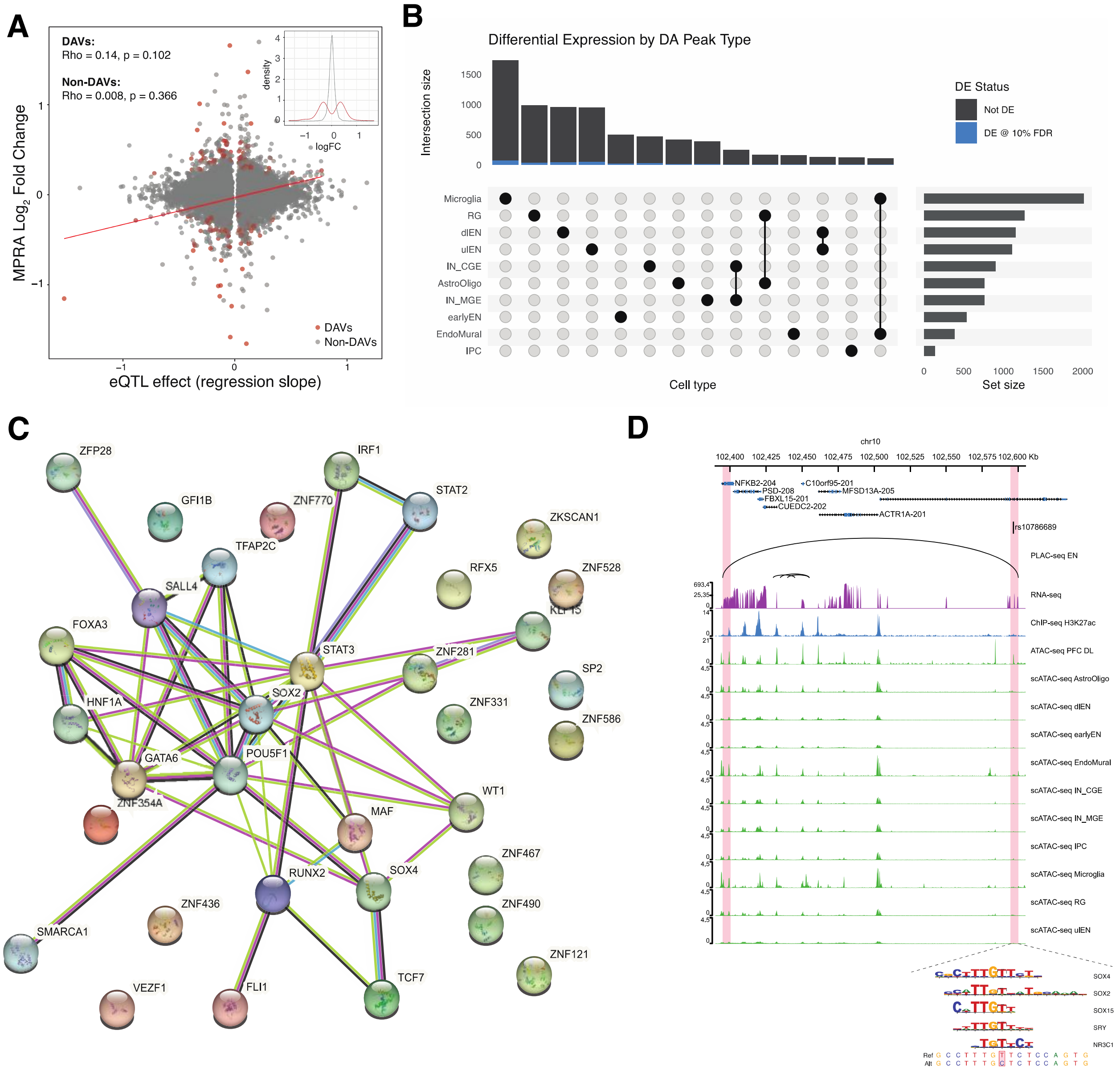


**Fig. S5. lentiMPRA identifies potential causal regulatory variants influencing risk genes of psychiatric disorders.** (**A**) Scatter plot showing the MPRA $\log_2$ Fold Change (y-axis) versus the regression slope of PEC eQTL (x-axis). Red dots indicate DAVs, the red line is its linear regression line, and rho is Pearson's correlation. Inset shows the density distribution of $\log_2$ Fold Change in DAV (red) or non-DAV (gray). (**B**). Upset plot showing the number of variants (bar) located in cell-type-specific DA regions (dots and lines below). The number of DAVs is highlighted in blue. (**C**) Protein-protein interaction (PPI) network using STRING (*73*) for TFs disrupted by DAVs. (**D**) Genome browser tracks showing an SCZ-related regulatory variant rs10786689, that was predicted to increase *NFKB2* expression via disrupting binding sites of SOX4/SOX2/SOX15/SRY/NR3C1. The top track shows a PLAC-seq chromatin loop in EN (*32*), the second track shows bulk RNA-seq in primary cortical cells, the third track shows bulk H3K27ac ChIP-seq (*31*), followed by a track of bulk ATAC-seq in the deep-layer cortex (*31*). The bottom ten tracks show scATAC-seq from the developing human cortex (*11*).

**Table S1. Characterization of active cell-type-specific DA regions.** For each cell type, active DAs showed higher means across several features, compared with inactive ones. This table lists q-values from the Wilcoxon test. Blank cells indicate that no chromatin interaction data was available for identifying gene targets of DAs. This data is represented in Figure 2C.

| cell_type | Conservation | Target Expression | # Motifs | # USF Motifs |
|---|---|---|---|---|
| AstroOligo | 2.70E-02 | | 4.98E-09 | 5.18E-10 |
| dlEN | 5.95E-08 | 1.64E-04 | 7.02E-01 | 5.27E-01 |
| earlyEN | 2.56E-04 | 1.11E-01 | 9.27E-01 | 6.27E-01 |
| EndoMural | 2.88E-02 | | 1.81E-04 | 3.05E-05 |
| IN_CGE | 1.40E-14 | 4.41E-01 | 1.12E-01 | 5.02E-03 |
| IN_MGE | 2.29E-09 | 4.00E-03 | 9.20E-01 | 6.08E-01 |
| IPC | 2.47E-01 | 3.65E-01 | 1.64E-01 | 1.05E-01 |
| Microglia | 5.52E-03 | | 2.91E-03 | 8.86E-07 |
| RG | 3.63E-05 | 8.45E-02 | 2.31E-03 | 3.03E-04 |
| ulEN | 1.66E-03 | 3.72E-02 | 9.99E-01 | 6.87E-01 |

**Table S2. DA regions validated for cell-type-specific enhancer activity.** Eleven DA regions specific to different cortical cell types were tested individually via reporter assays to confirm their regulatory function and cell-type specificity.

| Name | Genomic coordinate (hg38) | Cell type (scATAC-seq) | MPRA ratio | Age | Enhancer | Cell specificity |
|---|---|---|---|---|---|---|
| EN-1 | chr2:165141999-165142269 | pan-EN | 2.68 | GW20 | YES | YES |
| uIEN-1 | chr2:152408730-152409000 | uIEN | 2.73 | GW22/20 | YES | NO |
| uIEN-2 | chr5:89274678-89274948 | uIEN + earlyEN | 2.47 | GW19 | YES | YES |
| dIEN-1 | chr14:70993483-70993753 | dIEN | 2.46 | GW16 | YES | NO |
| dIEN-2 | chr9:14297525-14297795 | dIEN | 2.45 | GW16 | YES | YES |
| Astro/Oligo-1 | chr5:43217050-43217320 | Astro/Oligo | 2.26 | GW20 | YES | NO |
| Astro/Oligo-2 | chr14:50985465-50985735 | Astro/Oligo | 2.13 | GW20 | YES | NO |
| RG-1 | chr6:19663461-19663731 | RG + IN | 2.15 | GW16 | YES | NO |
| RG-1 | chr21:16154321-16154591 | RG + Astro/Oligo | 2.1 | GW16 | YES | NO |
| tRG-1 | chr1:68141016-68141286 | RG + Astro/Oligo | 1.56 | GW18 | YES | NO |
| tRG-2 | chr22:43331407-43331677 | RG | 1.45 | GW18 | YES | NO |

**Data S1. (separate file)**

S1. DA-library-ratios.tsv

S2. Variant-library-ratios.tsv

S3. de-rna-primary_vs_organoid.tsv
